# Cell type specific allometry controls sex-differences in *Drosophila* body size

**DOI:** 10.1101/2025.08.25.671808

**Authors:** Soumitra Pal, Jerome Avellaneda, Celena M. Cherian, Puja Biswas, Georg Vogler, Elizabeth J. Rideout, Frank Schnorrer, Teresa M. Przytycka, Brian Oliver

## Abstract

Species and sex-specific differences in organ size are fundamental features of animal biology, yet the mechanisms that drive these differences remain debated. Adult female *Drosophila* are larger than males. While most organs are present across both sexes, the underlying mechanisms driving sex-specific organ and body size scaling of *Drosophila* remain unclear. Using single-nucleus transcriptomes from the Fly Cell Atlas, combined with experimental validation, we show that different *Drosophila* organs scale through distinct strategies, including cell size, cell number, or a combination of both, in an allometric rather than uniform manner. Larger female flight muscles develop from more myoblasts than in males, while cardiomyocyte numbers are the same despite forming a larger heart in females. Female fat body cells are larger and express more ribosomal protein-coding mRNAs, supporting increased cell size. In contrast, males have a greater number of fat body cells. Together, this sex-specific allometry in cell size and number define the cellular basis for differences in body and organ size between sexes in *Drosophila*. By uncovering how a conserved developmental system produces sex-specific proportions through distinct cellular strategies, our work offers a framework for dissecting sex differences in other species and systems.

## INTRODUCTION

Organ morphogenesis and homeostasis must be meticulously coordinated to ensure the establishment of a functional body plan in adulthood. Disruptions in proportionality,whether due to undergrowth or overgrowth, can lead to reduced fitness or diseases such as cancer. Despite extensive research into growth and developmental pathways, how proportionality across organs and tissues is regulated remains incompletely understood ^1^.

A key, yet underexplored, dimension of proportionality is sex. In many animals, males and females follow distinct allometric trajectories, producing sex-specific body plans. These dimorphic traits are not just curiosities of evolution; they offer a natural experiment in developmental scaling. Understanding how male and female body plans are differentially optimized can provide fundamental insights into the mechanisms that govern proportional growth.

Sex is a fundamental biological variable that influences nearly all aspects of development, physiology, and disease susceptibility. However, our understanding of sex-based differences still lags far behind our knowledge of general developmental and morphological processes. One of the most widespread across animal kingdom yet poorly understood manifestations of this gap is sexual size dimorphism (SSD), where individuals of one sex are consistently larger than the other ^2–8^. In *Drosophila melanogaster*, adult females are substantially larger than males (Fig. 1A1), yet both sexes develop from genetically similar zygotes and share most organs and cell types.

**Fig. 1.**
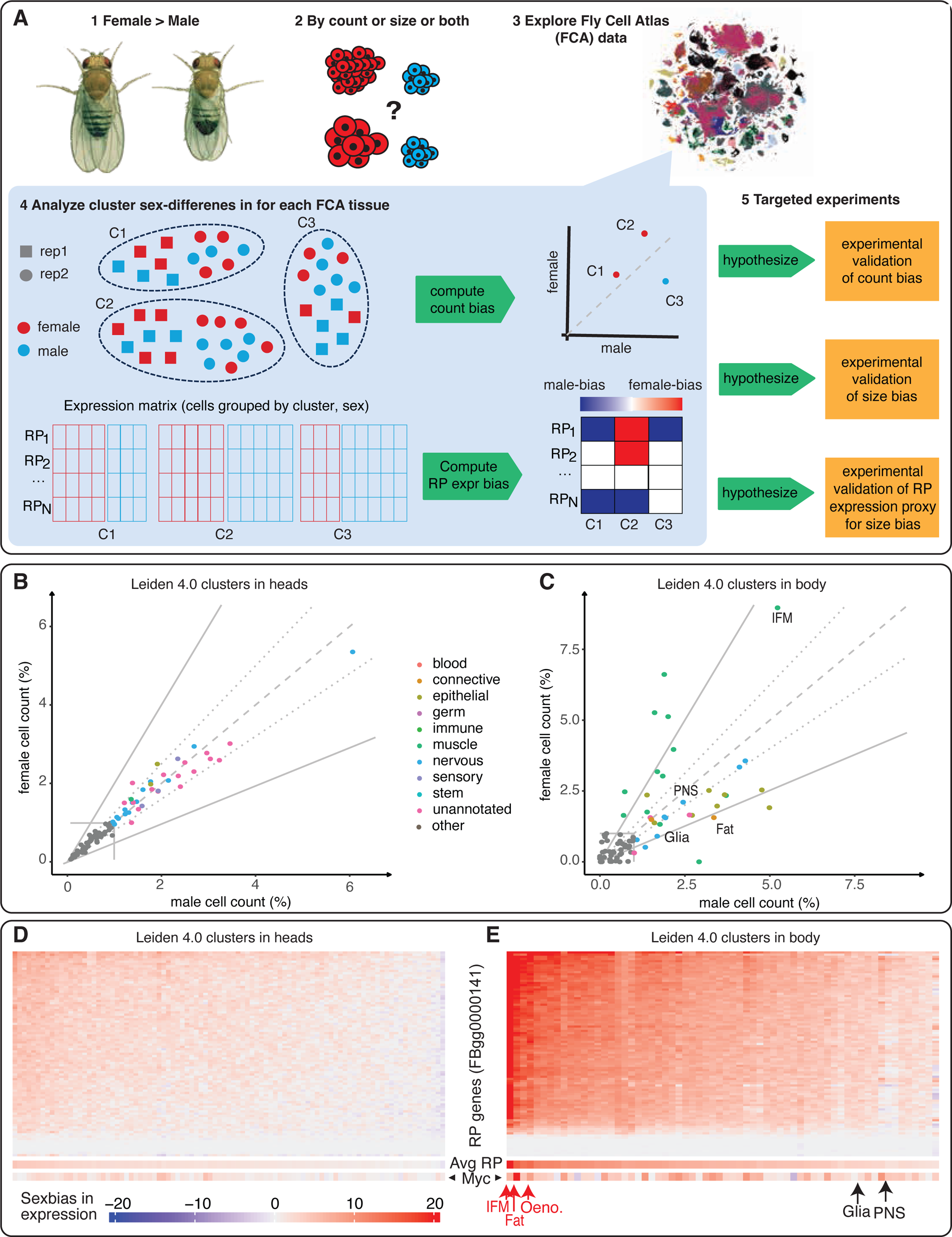
Overview of the study, highlighting sex-specific differences in cell number and size in the single-nucleus Fly Cell Atlas (FCA). **(A)** Sex-related phenotypic differences, including variations in size (1), may be attributed to differences in cell type nuclei proportions, cell size, and/or gene expression (2). We utilize the single-nucleus Fly Cell Atlas (FCA), which includes 15 individually dissected, sex-specific tissues, along with whole-head and headless-body datasets. These are visualized as t-SNE plots, colored by annotated cell type clusters (3). In this analysis, sex-specific clusters (black in tSNE plot) are excluded. The head and headless body tissues were sequenced in four biological replicates each. These samples, free from potential dissection-related variability, were used to assess sex-related biases in cell counts (4 top). Cluster-specific ribosomal gene expression biases suggest and are used as a proxy for potential differences in cell sizes (4 bottom). Differences in cell type proportions and sizes were experimentally validated (5). (**B, C)** Scatter plots of the FCA Leiden 4.0 clusters for head samples (**B**) and headless body samples (**C**). The Y-axis represents the percentage of female cells, and the X-axis represents the percentage of male cells, both normalized by the total cell count in the replicated samples. The dashed line indicates equal proportions of female and male cells, while the dotted lines show a cell proportion bias of 1.25-fold, and the solid lines represent a bias of 2.0-fold. Female-biased clusters are shown in red, and male-biased clusters in blue, with darker shades representing a bias of at least 2.0-fold. Clusters containing more than 1% of cells are color-coded based on their nine broad annotations, while those below this threshold are shown in gray. (**D, E**) Heatmaps displaying the expression bias of ribosomal protein genes (RP; FBgg000141) genes (rows) across Leiden 4.0 clusters (columns) in the head (D) and headless body (**E**) samples. Both heatmaps use a consistent color scale. At the bottom of each heatmap, the average expression bias of all RP genes as well as the expression bias of *Myc* are also shown.

The origins of SSD can, in principle, arise from differences in (i) gene expression, (ii) the presence of sex-specific cell types, (iii) the number of cells, or (iv) the size of those cells. Previous research in *Drosophila* has largely focused on gene expression in sex-specific organs like the gonads ^9–12^, which are governed by a well-characterized sex-determination pathway ^13–15^ However, whether and how scaling differences in shared, non-sex-specific tissues are achieved via changes in cell size and number remains largely unexamined (Fig. 1A2).

Recent advances in single-cell transcriptomics allow for high-resolution mapping of cell type abundance and characteristics across complex tissues. The single-nucleus Fly Cell Atlas (FCA) ^16^ provides such a dataset, profiling over 580,000 nuclei from 15 anatomically dissected, sexed tissues, as well as whole head and body samples from both sexes (Fig. 1A3). This unique resource enables a systematic investigation of SSD at the cellular level. By excluding gonadal cells and focusing on shared cell types, we quantified sex differences in the number and size of nuclei for each cell type (Fig. 1A4). Here, we aim to address two core questions: 1) Are sex differences in organ and body size driven primarily by changes in cell number, cell size, or both? and 2) Do these differences apply broadly across shared cell types or are they restricted to specific tissues? We then validated several findings experimentally in muscle, heart, and fat body cells—three metabolically and functionally distinct tissues—to confirm that sex-specific scaling arises from distinct combinations of increased cell size, increased cell number, or both (Fig. 1A5).

This study offers a comprehensive, cellular-resolution analysis of sexual size dimorphism in a model organism, uncovering how differences in cell number and size contribute to sex-specific body plans. These findings have broad implications for understanding growth control, sex-biased disease susceptibility, and evolutionary developmental biology. By leveraging a whole-body single-nucleus atlas, we illuminate a fundamental, yet understudied, aspect of organismal biology: how sex shapes size at the level of individual cell types.

## RESULTS

### Non-uniform allometric scaling revealed by computational analysis of Fly Single Cell Atlas

For the analysis of sex-differences in cell number and ribosomal content in the FCA, we focused on the head and body datasets (Table S1), which include replicated samples across all adult tissues. While most non-sex-specific cell types exhibited similar nuclear counts between sexes (Fig. 1B-C), there were significant sex-specific differences in the proportions of male and female nuclei across several cell types (Fig. 1C; Figs. S1-S2 for unreplicated dissected tissues). For example, multiple muscle cell types, including the large indirect flight muscles in the thorax, had a pronounced female bias (Fig. 1C). This observation aligns with findings in mammals, in which muscle nuclear number scales with the muscle volume ^17–19^. In contrast, cell types with a higher proportion of male nuclei included epithelial, glial, and perineurial glial sheath cells (Fig. 1C). Interestingly, females had a lower proportion of fat body cell (adipocytes) nuclei than males (Fig. 1C). Notably, sex differences in cell numbers were more pronounced in the headless body samples (Fig. 1C) than in head samples (Fig. 1B), indicating that non-uniform scaling patterns vary by major body part.

Ribosome abundance is known to scale with cell size ^20–22^. We therefore used ribosomal protein-encoding gene (RP) expression bias as a proxy for sex differences in cell size (Tables S2-S3; Fig. 1A4, D-E for head and body; Figs. S3-S16 for dissected tissues). Because ribosomes are multi-subunit complexes with strict stoichiometry ^23^, we first tested whether sex-biased RP expression was stoichiometric, i.e., correlated across RP genes. Indeed, we observed strong correlations in sex-biased expression of RP genes across cell types (Fig. S17A-B). Further supporting this, Myc, a key regulator of ribosome biogenesis ^24^, also exhibited sex-biased expression patterns consistent with RP expression bias (Fig. 1D-E, bottom of heat map). These findings, including those for several transcription-related genes (Fig. S17C-D), indicate that ribosomal gene expression bias is systematic and serves as a reasonable proxy for sex differences in cell size.

The magnitude of the sex bias in RP gene expression varied across cell types (Fig. 1D-E). For example, strong female-biased RP expression was observed in indirect flight muscles, oenocytes, and the adult fat body (Fig. 1E), whereas glia and neurons showed only a slight female bias. Notably, despite having a male-biased cell number, the fat body cells exhibited markedly higher female-biased RP gene expression (Fig. 1E), suggesting that the lower cell number in females is offset by increased cell size.

Overall, these data indicate that non-uniform sex differences in cell number exist across multiple cell types, some overrepresented in females, others in males. Molecularly, sex-differences in RP gene expression, and thus cell size, is also non-uniform.

### Larger female indirect flight muscles contain more nuclei

To directly demonstrate the striking female bias in flight muscle nuclei number observed in the FCA data, we quantified adult flight muscle size (Methods; Tables S4-S5). Indirect flight muscle cells are syncytia containing thousands of nuclei built by myoblast fusion during development ^25,26^. Adult flies have 12 dorsal longitudinal flight muscle cells (DLMs) which span the thorax from anterior to posterior ^27^. DLMs are longer in females than in males, commensurate with the longer females thoraxes ^2^. To precisely quantify total DLM volume in both sexes, we measured DLM muscle length in longitudinal sections (Fig. 2A, B) and cross-sectional area in 3-to-5-day old adult males and females (Fig. 2C, D). Using these measurements, we determined that the total volume of the 12 DLMs in females is ∼50% greater than in males (Fig. 2E).

**Fig. 2.**
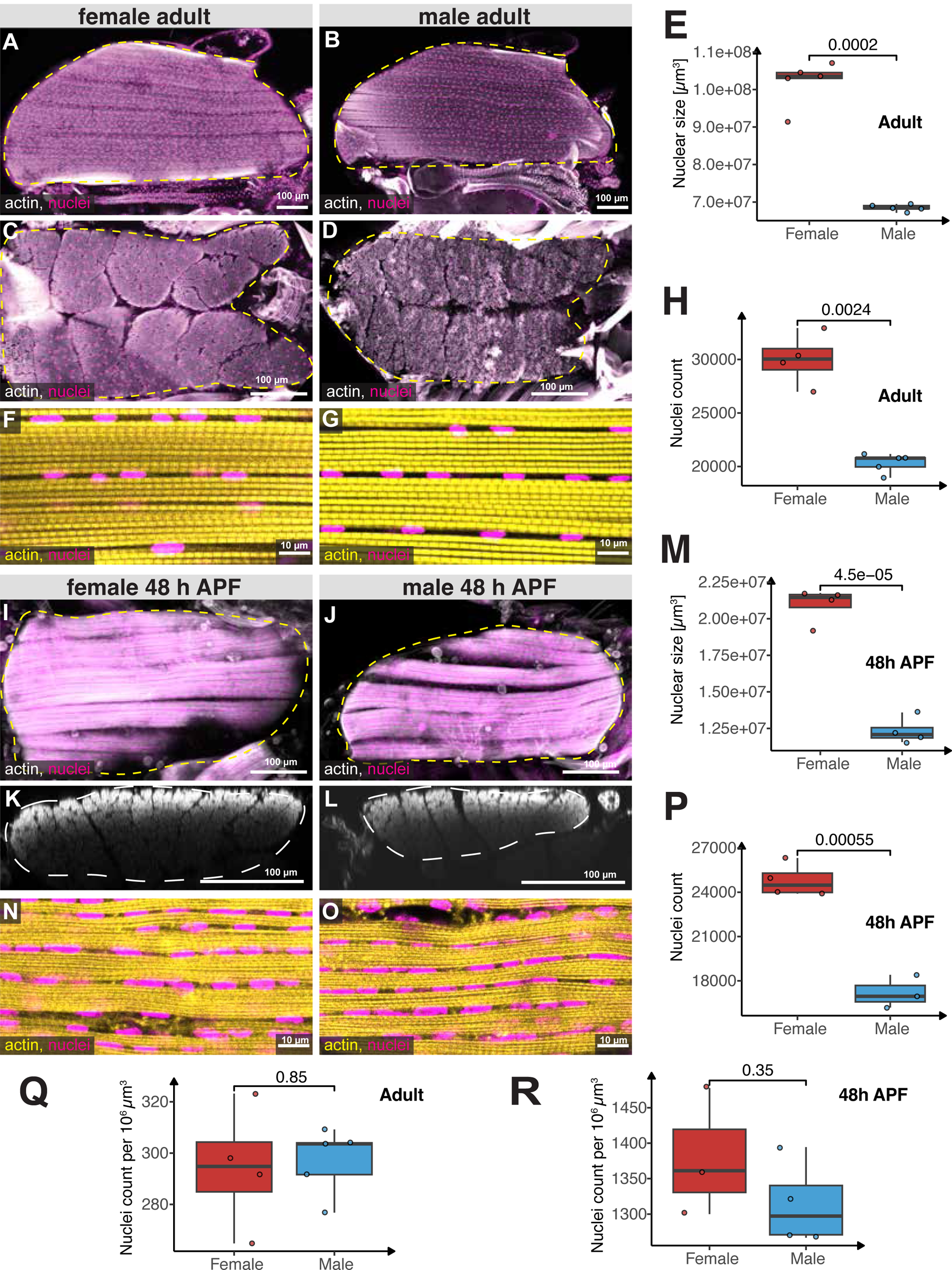
Sexual allometry of the indirect flight muscles. (**A-D**) Hemi-thoraces (**A-B**) of adult females (**A, C**) or males (**B, D**) expressing histone-RFP (*Mef2-GAL4, UAS-histone-RFP*) in all muscles to label nuclei (magenta) were cut longitudinally (A, B), or sagittally (cross-section) (**C-D**) and stained with phalloidin to label actin (gray). The yellow dotted lines mark the outline of the dorsal longitudinal flight muscles (DLMs); note that female flight muscles are larger compared to male muscles. (**E**) Quantification of the 12 DLMs volume in adult females (red) and males (blue). (**F-G**) High magnification of adult flight muscles showing actin (yellow) and nuclei (*Mef2-GAL4, UAS-histone-RFP*; magenta). (**H**) Quantification of the nuclei number in the 12 DLMs of adult females (red) and males (blue). (**I-L**) Developing flight muscles at 48 h APF of females (**I, K**) or males (**J, L**) displaying actin (gray) and nuclei (magenta) by longitudinal view (**I, J**) and cross-section view, reconstructed as xz-projections (**K, L**). (**M**) Quantification of muscle size of the 12 developing DLMs at 48 h APF for females (red) and males (blue). Note developing DLMs are larger in females compared to males. (**N-O**) High magnification of longitudinal sections of developing flight muscle at 48 h APF showing actin (yellow) and nuclei (magenta). (**P**) Quantification of nuclei numbers of the 12 developing DLMs at 48 h APF for females (red) and males (blue). Note female muscles contain more nuclei. (**Q-R**) Quantification of muscle nuclei density in adult (**Q**) and developing (**R**) flight muscles in females (red) and males (blue). Note that nuclear density in males and females is comparable and decreases during development when the muscles grow in size. Scale bars are 100 µm for (**A-D**, **I-L**) and 10 µm for (**F, G, N, O**). Each box plot (**E, H, M, P-R**) displays the first and third quartiles as the hinges, with the median represented by the line in the middle. The upper whisker extends from the upper hinge to the largest value within 1.5 times the interquartile range (IQR) from the hinge (IQR is the distance between the first and third quartiles). Similarly, the lower whisker extends from the lower hinge to the smallest value within 1.5 times the IQR. *p*-values are calculated using the t-test.

To quantify the number of nuclei in flight muscles, we expressed histone-RFP in all indirect flight muscles and counted the DLM nuclei in a representative volume (Fig. 2F, G, see Methods). We found approximately 20,000 nuclei in male DLMs and 30,000 nuclei in female DLMs (Fig. 2H), consistent with ∼1.5-fold female-bias in the FCA data (Fig. 1F). Thus, the ∼50% larger female flight muscles contain ∼50% more nuclei, resulting in comparable nuclear densities (Fig. 2Q). This indicates that larger female muscles recruit more myoblasts during development.

Muscle fibers develop through myoblast fusion between 6 and 24 hours after puparium formation (APF) ^26,28^. To determine whether the sex difference in size and nuclei number had occurred by pupal stages, we dissected and stained developing DLMs at 48 hours APF, a stage when myoblast fusion is complete. We imaged the entire volume of the 6 DLMs in each hemithorax of pupae, using phalloidin to visualize actin and measure the fiber length and cross-sectional area (Fig. 2I-M). As expected, the muscle fiber volume at 48 hours APF is about five times smaller than in adults. However, female muscles are already about ∼50% larger than male muscles at this stage, indicating that the sex-specific cell size difference is established early in muscle development.

To quantify nuclei numbers post-myoblast fusion, we counted histone-RFP nuclei within a representative volume of male and female DLMs (Fig. 2N-O). The sex-specific difference in nuclei number observed at 48 hours APF mirrored the difference found in adults (Fig. 2P-R), demonstrating that the sexually dimorphic flight muscle nuclear numbers result from increased myoblast fusion during development. Given that only 2,000–4,000 myoblasts are present in third instar larvae ^29,30^, this dimorphism likely emerges during early pupal stages. Thus, the larger female flight muscles are constructed by a larger number of myoblasts.

### Larger female hearts have larger cardiomyocytes

Not all muscle types exhibited differences in nuclei number. In *Drosophila*, the heart forms a single contractile tube composed of contractile cardiomyocytes (CMs) and morphologically distinct inflow tracts, including segmentally repeating ostia and valve cells that regulate cardiac hemodynamics ^31^. The adult heart consists of 84 CMs, compartmentalized in four contractile chambers, with no sexual dimorphism in CM number. However, we observed strong sex differences in heart chamber size and contractile performance (Methods; Tables S4, S6). Female hearts exhibit larger end-diastolic (EDD; Fig. 3A-B) and end-systolic diameters (ESD; Fig. S18A), leading to a 30% increase in stroke volume (SV; Fig. 3B’). This suggests that larger females pump more blood with the same number of cardiac cells. Interestingly, heart contractility (fractional shortening, FS) remains identical between sexes (Fig. 3B”). Thus, unlike the indirect flight muscle, the sex-specific physiology of the heart is associated with increased organ size without changes in cell or nuclear number in females.

**Fig. 3.**
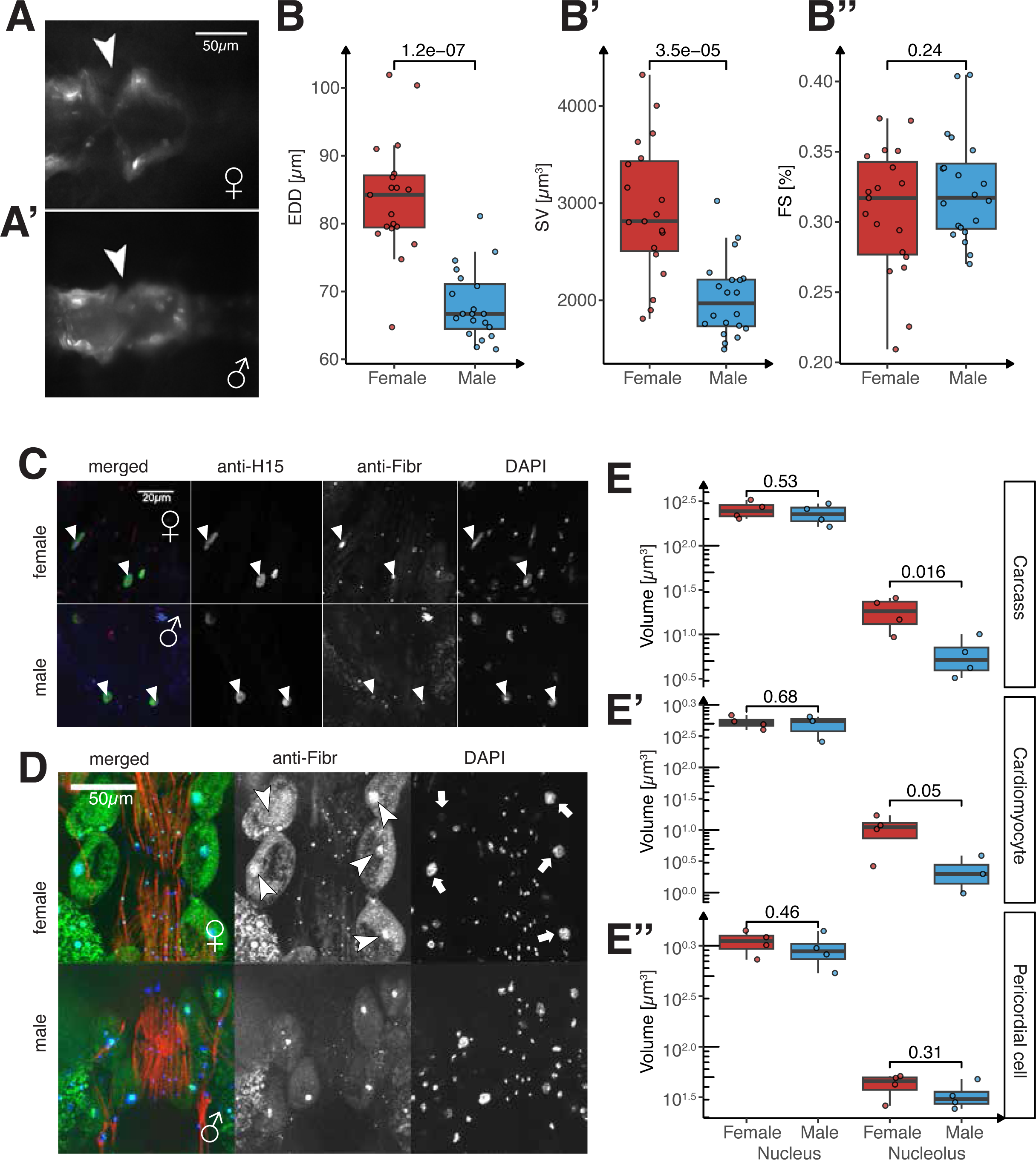
Sexual allometry of the fly heart. (**A-B**) tdTomato-labeled female (**A**) and male (**A’**) heart chambers from segment A2/3. Inflow tract (ostia) indicated (arrowheads). Sex differences are reflected in heart size (end-diastolic diameter, EDD; **B**) and stroke volume (SV; **B’**) but not contractility (FS; **B’’**). (**C**) Adult cardiomyocyte nuclei (green, anti-H15) were labeled with nucleolar marker Fibrillarin (red) and DAPI (blue) for quantification of nuclear and nucleolar sizes. (**D**) Heart-associated pericardial cells (PCs, arrowheads) are morphologically distinct with large nucleoli (arrowheads) inside large nuclei (arrows). PCs are localized along the heart and ventral layer (myofibrils, red). (**E**) Quantification of nuclear (DAPI, blue) and nucleolar (Fibrillarin, green) volume in female (red) and male (blue) nuclei. Measurements are from carcass (including fat body; **E**), cardiomyocyte (**E’**), and pericardial cell nuclei (**E’’**). Each box plot (**B, E**) displays the first and third quartiles as the hinges, with the median represented by the line in the middle. The upper whisker extends from the upper hinge to the largest value within 1.5 times the interquartile range (IQR) from the hinge (IQR is the distance between the first and third quartiles). Similarly, the lower whisker extends from the lower hinge to the smallest value within 1.5 times the IQR. *p*-values are calculated using the t-test.

To understand how the same number of cells can generate a larger heart in females (Fig. S18B), we analyzed the sex-differences in RP expression the FCA, which suggested that the cells are indeed larger (Figs. 1F-G, S18C). This led us to investigate whether increased female ribosomal protein gene expression correlated with differences in nuclear and nucleolar sizes, as the nucleolus is responsible for producing and assembling ribosomal RNAs into ribonuclear complexes ^32^.

The presence of multiple cell types in and around the heart allowed us to observe and quantify nuclear and nucleolar sizes in these cell types, particularly CMs, and pericardial cells (PCs). Using antibody staining for the nucleolar protein Fibrillarin ^33,34^, we found that female nucleoli are larger, both in absolute terms and relative to nuclear volume, in CMs and PCs (Fig. 3C-E). Interestingly, total nuclei volume in CMs was not different between sexes (Fig 3E), suggesting that ploidy differences which are common in *Drosophila* ^35–37^, do not play a role in generating a larger female heart.

Thus, our findings indicate that female heart growth is not driven by additional cell recruitment but rather by increased ribosome production, and increased cell size. Interestingly, the postmitotic growth mechanism also drives mammalian heart size ^38^.

### Female bias in ribosomal gene expression in fat body cells leads to sex biased protein synthesis

The fat body is an endocrine and energy-storing organ that is functionally equivalent to mammalian adipose tissue, and is composed of large polyploid cells ^39^. The FCA data indicate a profound female bias in RP mRNA in the adult fat body (Fig. 1E), which we confirm is also present in the larval fat body (Fig. 4A; Methods; Tables S4, S7; Fig. S17E for adults). Also aligning with FCA data is our finding that while females show more fat body cells overall (Fig. 4B), male larvae have proportionally more fat body cells when their smaller body size is taken into account (Fig. 4C). Since ribosomes scale with cell size, one potential explanation for this female bias in RP gene mRNA levels is a sex difference in cell size. Indeed, one prior study showed that larval fat body cells are larger in females than in males ^3^. To test whether a sex difference in larval cell size is the sole determinant of the sex bias in RP gene expression, we measured nuclear and nucleolar size in fat body cells from both males and females (Fig. S19A). We found that female cells had larger nuclei than males, consistent with their larger size. Female fat body cells also had larger nucleoli (Fig. 4D) the primary site for ribosome biogenesis. While this larger nucleolar size aligns with a larger nuclear and cell size, the female bias in nucleolar size persists even after normalization for nuclear size (Fig. 4E). This suggests in larvae, female cells have proportionally more ribosome biogenesis even after cell size is taken into account.

**Fig. 4.**
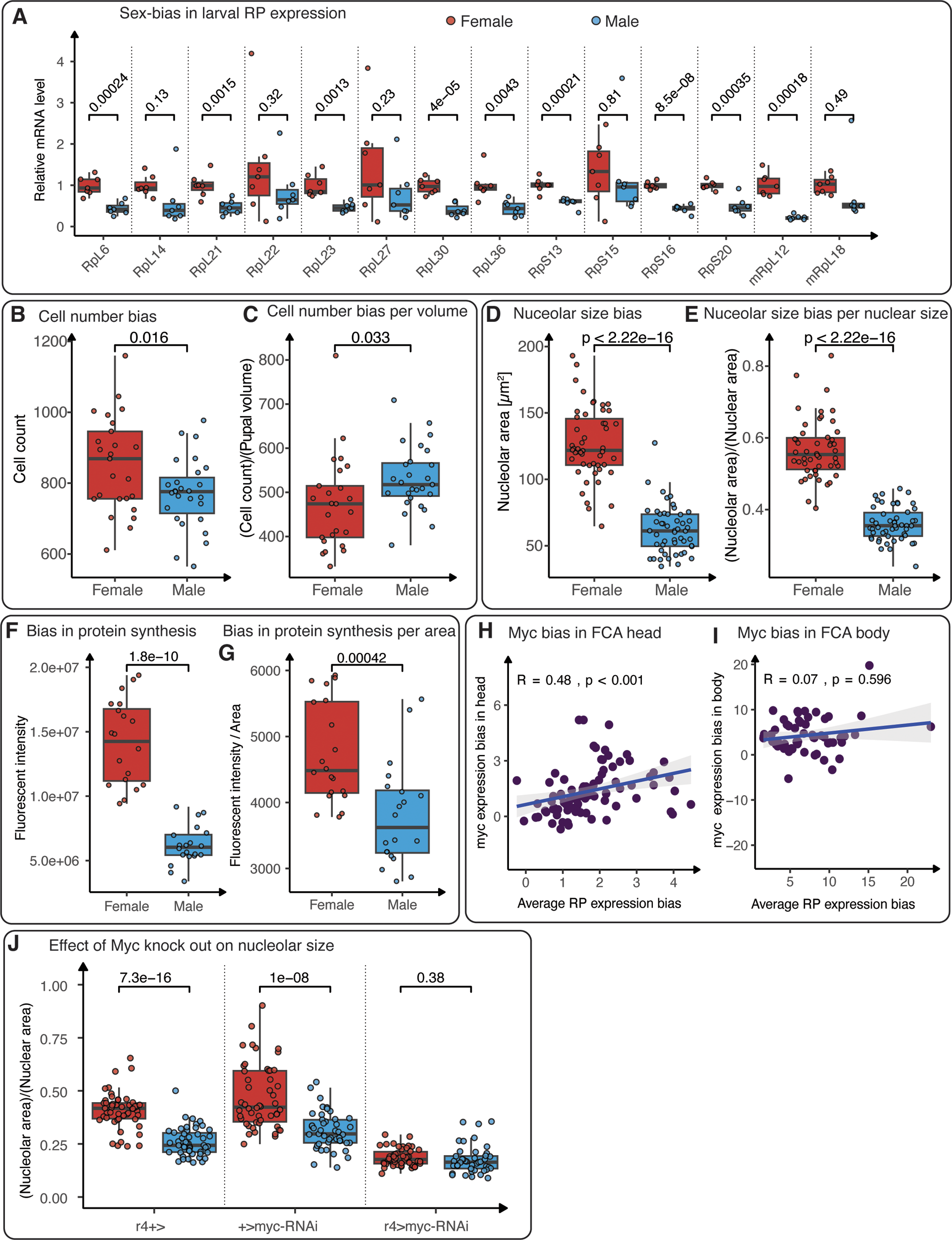
Sexual allometry of the fat body and ribosomal protein expression as a proxy for cell size. (**A**) mRNA levels of large (RpL), small (RpS), and mitochondrial large (mRpL) ribosomal subunit genes in *w*^1118^ female (red) and male (blue) adult abdominal carcasses demonstrate higher RP gene expression in females, with the difference being significant in most cases. (**B**) Quantification shows that the number of larval fat cells is significantly higher in females (red) compared to males (blue) at 108 hr AEL. (**C**) However, when the number of larval fat cells is normalized to body volume, males exhibit a higher cell count. (**D**) Quantification of the nucleolar area shows that female (red) nucleoli are significantly larger than male (blue) nucleoli in the larval fat body. (**E**) The nucleolar-to-nuclear area ratio is also significantly higher in the female fat body compared to the male fat body. (**F**) Quantification of total fluorescence intensity showed that female fat cells have significantly higher levels of protein synthesis than male fat cells. (**G**) Protein synthesis levels remain higher in females, even when total fluorescence intensity is normalized for sex differences in cell size. (**H, I**) Scatter plots showing the expression bias of *Myc* gene relative to the average expression bias of RP genes in Leiden 4.0 clusters for the head (**H**) and headless body (**I**) samples. Spearman’s correlation coefficients and *p*-values are indicated. (**J**) Fat body knockdown of *Myc* caused a significant decrease in nucleolar size in both sexes (GAL4 and UAS controls vs. *Myc* fat body knockdown) and abolished the sex difference in nucleolus size (female vs. male *Myc* fat body knockdown). Each box plot (**A-G, J**) displays the first and third quartiles as the hinges, with the median represented by the line in the middle. The upper whisker extends from the upper hinge to the largest value within 1.5 times the interquartile range (IQR) from the hinge (IQR is the distance between the first and third quartiles). Similarly, the lower whisker extends from the lower hinge to the smallest value within 1.5 times the IQR. *p*-values are calculated using the t-test.

Given that ribosome biogenesis is linked with cellular protein synthesis, we next asked whether female fat body cells synthesize more proteins than males. We monitored nascent protein synthesis in female and male larval fat bodies ^40^. We found that nascent protein synthesis was significantly higher in female fat body cells than in males (Fig. 4F). This female-biased protein synthesis could not be solely attributed to larger cell size in females, as protein synthesis remained higher in females even after we normalized for cell size (Fig. 4G). Thus, female fat body cells have proportionally more ribosome biogenesis and protein synthesis than male fat body cells when cell size is taken into account. This increased biosynthetic activity in fat body cells may contribute to cell size differences, but also to the regulation of body size via production of factors that mediate body growth via interorgan communication ^41–45^.

We next wanted to identify factors that might contribute to the sex difference in RP expression in the larval fat body. Because *Myc*, has a strong female bias in gene expression (Fig. 1D-E; 4H-I), we used fat body driver *r4-GAL4* to knock down *Myc* mRNA levels. We found that loss of Myc significantly reduced nucleolar size in both sexes, eliminating the sex difference in nucleolar size (Fig. 4J). This demonstrates that Myc plays a key role in regulating the sex difference in nucleolar size.

## DISCUSSION

Our work demonstrates that both tissue-specific differences in cell number and cell size drive sexual size dimorphism in *Drosophila* and non-uniform allometric scaling across the organism. Female-biased nuclear counts in indirect flight muscles, male-biased counts in fat body cells, enhanced ribosomal protein (RP) gene expression in female fat body and heart tissues, jointly indicate that different tissues employ distinct developmental strategies to achieve sex-specific sizes. These findings shed light on sexual size differences that are widespread across animals ^46^, including humans, in which males are generally larger. Such differences likely result from the optimization of total body size and organ allometry between sexes, within the constraints of a shared genome ^47^.

### Tissue-specific mechanisms of sexual size dimorphism

The female-biased nuclear counts in indirect flight muscles, established during pupal myoblast fusion, point to sex-specific regulation of myogenesis. The ∼50% larger female dorsal longitudinal muscles (DLMs) contain proportionally more nuclei, maintaining comparable nuclear density to males, mirroring mammalian muscle progenitor recruitment modulated by sex hormones like testosterone ^17–19^. To rule out developmental timing as a confounder, we confirmed that nuclear differences persist from 48 hours after puparium formation (APF) to adulthood, indicating that dimorphism stems from cell-autonomous sex identity. These findings position Drosophila as a valuable model for dissecting developmental mechanisms underlying sex-specific muscle growth, with potential relevance to mammalian myopathies.

In contrast, the larger female heart achieves increased size without additional cardiomyocytes, driven by enhanced ribosome biogenesis and larger cell sizes. This resembles mammalian cardiac hypertrophy, where ribosome-driven growth supports increased heart size in response to physiological demands, such as pregnancy or exercise ^38^. The absence of ploidy differences in female cardiomyocytes suggests that ribosome biogenesis, rather than nuclear content, drives heart size dimorphism, ensuring functional equivalence between sexes, as evidenced by preserved heart contractility despite increased stroke volume in females.

### Cell-autonomous and hormonal regulation

Organ size regulation is critical for normal development and physiology across organisms, involving both intrinsic cell-autonomous processes and extrinsic hormonal controls ^1^. In *Drosophila*, coordinated growth during metamorphosis is a textbook example of hormonal control ^48^, while cell-autonomous mechanisms are less commonly highlighted. The cell-autonomous component of sex-biased body size is evident in mixed-sex gynandromorphs, where XX female portions exhibit larger abdomens, wings, and eyes compared to X0 male portions ^49^. This cell autonomous effect may be mediated by X chromosome number, with our findings showing that the X-linked Myc gene is cell-autonomously required for fat body cell size and sexual dimorphic size. While Myc has been suggested to escape dosage compensation ^50^, our data reveal dynamic Myc expression rather than a simple two-fold increase in female cells, suggesting a complex interplay with female-specific genetic regulatory networks, such as Sex-lethal and transformer ^3,7,68^. These genes, broadly expressed across cell types ^51^, may drive the widespread upregulation of RP genes in females, contributing to sex-specific allometry.

### Limitations and future directions

Despite the robustness of our findings, limitations remain. Direct cell volume measurements across all FCA cell types were not feasible due to technical constraints, however RP expression served as a reasonable proxy for cell size. Future studies could leverage advanced imaging to confirm these differences. Additionally, functional assays beyond heart stroke volume, such as flight muscle endurance or fat body lipid storage, could clarify the physiological impacts of sex-specific cell size and number. The lack of *Myc* rescue experiments limits mechanistic conclusions, but temporal overexpression studies could address this. Analyzing earlier developmental stages, such as larval myoblast pools, could further confirm the onset of dimorphism and exclude developmental timing effects.

Future studies could investigate molecular signals, such as sex hormones or insulin-like peptides, driving differential myoblast fusion or ribosome biogenesis, potentially revealing how these mechanisms coordinate systemic growth via interorgan signaling. Comparative studies in mammals could test whether Myc-driven ribosome biogenesis underlies sex differences in muscle or adipose tissues, informing conditions like sarcopenia or cardiac hypertrophy.

## Conclusion

Our findings highlight the power of single-cell atlases to uncover tissue-specific sexual dimorphism, offering a framework for studying allometric scaling across organisms. The interplay between cell-autonomous regulation, driven by genes like *Myc*, and genetic pathways, including Sex-lethal and transformer, underscores the complexity of sexual dimorphic size development. These mechanisms, conserved across species, have broader implications for understanding development and diseases, such as cancers involving the *Myc* oncogene.

## MATERIALS AND METHODS

### Analysis of sex-difference in Fly Cell Atlas (FCA)

#### Datasets

Loom files for each of the 14 (non-sex-specific) tissue datasets were downloaded from FCA (https://flycellatlas.org) website. The details of these datasets are provided in the sheet named “FCA_Samples” in Table S1. The loom files were converted to AnnData format using loompy v3.0.7 (https://linnarssonlab.org/loompy) and anndata v0.10.9 ^52^ python packages.

#### Clustering Resolution

To address potential biases arising from the high number of unannotated nuclei in the FCA dataset, we employed Leiden clustering instead of relying solely on FCA-provided annotated cell types. For comprehensive analyses, Leiden clustering was applied at resolution 4.0 for the larger datasets (body and head) and resolution 1.0 for the dissected tissues. Clustering resolutions were selected based on manual inspection of the resulting number of clusters at each resolution and the distribution of male and female nuclei within each cluster (“Leiden_Clusters” in Table S1). The chosen resolutions ensured that the number of clusters was greater than the number of annotated clusters while avoiding clusters with very few cells. We annotated each of these Leiden clusters based on the predominant annotation of the cells within the cluster. These annotations were further grouped into nine broad classes: blood, connective, epithelial, germ, immune, muscle, nervous, sensory, and stem; and categorized by sex specificity: male-specific, female-specific, or non–sex-specific (see ‘FCA_Annotations’ in Table S1).

#### Filtering cells3

For our sex-difference analysis, we excluded all cells that were annotated as a sex-specific cell type. Additionally, we excluded cells that were annotated as “artefact” or had their sex labeled as “mix” in the FCA metadata.

#### Cluster Filtering

Clusters containing fewer than 100 male nuclei were excluded to avoid unreliable sex-differences inference.

#### Sex-bias in cell numbers

We compute sex-bias in cell numbers for the two replicated samples, *head* and *body*, each has 6 replicates, allowing for statistical significance.

For each cluster *c*, and for each replicate *r*, we calculate the normalized cell count for each sex *s* (female and male) as: 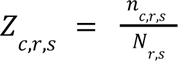, where *Z_c,r,s_* is the number of cells in cluster *c* for *c* replicate *r* of sex *s* and *N_r,s_* is the total number of cells for sex *s* in replicate *r*.

To assess sex bias, we compute the sex-biased cell count for each cluster and replicate as: 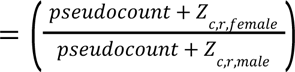 in which *pseudocount* is a small number such as 10^−256^.

The p-value for each cluster is computed using the Wilcoxon test on the values *Z*_*c*,*r,female*_ and *Z*_*c*,*r,female*_ for all *r*. Finally, Benjamini-Hochberg multiple testing correction is applied to the p-values for all clusters in each sample.

#### Normalization of expression

The raw UMI counts for all genes in each of the retained cells were normalized by dividing by the total number of UMIs across all genes in the cell and then multiplying by a factor of 10,000 using the `scanpy` function `sc.pp.normalize_total` with the parameter `target_sum=1e4`^53^, followed by log transformation (using the `scanpy` function `sc.pp.log1p`).

#### Sex-bias in expression

Sex-bias in gene expression was computed at two different resolutions: annotated clusters across all datasets, and additionally at Leiden resolution 4.0 for the head and (headless) body datasets. For each tissue and each cluster at the given resolution, the sex-bias of each gene was calculated as 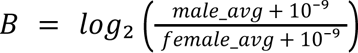, where *male_adg* and *female_avg* represent the average expression (calculated from the normalized expression matrix) of male and female cells, respectively, within the cluster. The p-value for the bias, adjusted for multiple testing, was determined using the Wilcoxon test. All these steps were performed using the `scanpy` function `sc.tl.rank_genes_groups`, with default parameters, to compare male and female cell groups within each cluster ^53^. Note that `rank_genes_groups` automatically un-logs when provided with the log-normalized expression.

For our analysis and visualization, we used the score generated by the `scanpy` function `sc.tl.rank_genes_groups.` The score reflects how consistently a gene’s expression is higher or lower in female cells compared to male cells. Specifically, we employ this score in the heatmaps for ribosomal protein (RP) and other related genes (see details in the section on sex-bias in size below) in Fig. 1D-E, S1A-D as well as the boxplots in Fig. S3B-C. To analyze the average sex-biased expression of RP genes, we compute their mean scores, which are visualized in Fig. 1D-E and in the scatter plots in Fig. 4H-I.

#### Sex-bias in size

We used the expression of ribosomal protein genes as a proxy for cell size. The sex-biased size of each cluster was computed as the average sex-biased expression of 94 genes from the FlyBase gene group FBgg0000141 (https://flybase.org/reports/FBgg0000141.htm; table S1), which includes 54 large ribosomal proteins (RpLs and few others) and 40 small ribosomal proteins (RpSs and few others). Additionally, we considered the sex-biased expression of 61 other genes known to be involved in translation and 9 specific genes of interest: DENR, InR, MCTS1, Myc, Nelf-A, NELF-B, Nelf-E, pix and tor (table S2).

### Experimental validation in muscle

#### Imaging of flight muscles

*Adults:* the head and abdomen from *Mef2-GAL4, UAS-histone-RFP* adult flies were removed with scissors. The remaining thoraxes were fixed for 30 min in PBS + 0.5% Triton (PBST) + 4% PFA on a nutator at room temperature (RT) and then washed 2x in PBST for 5 seconds and another time in PBST for 10 minutes. The fixed thoraxes were cut longitudinally or transversely using a microtome blade. The thorax halves were then transferred to glass dishes and stained for actin by incubation in phalloidin-Alexa 488 (1/500, Thermo Fisher Scientific #A12379) overnight at 4°C and then washed once rapidly with PBST and then 2x in PBST (10 min each) at RT. The thorax halves were mounted in Vectashield (SKU #H-1000-10) with 2 coverslips as spacer. To quantify nuclei number, actin (phalloidin-Alexa 488) and nuclei (histone-RFP) were imaged in a large z-stack with 40 µm depth on a Zeiss LSM880 confocal with 25x objective and images were processed with Fiji ^54^. To measure cell size, flight muscles were additionally imaged with a 10× objective to image the entire hemithorax in longitudinal and cross-sections.

*Pupae:* the sex of the pupa was determined visually by the presence of the testis. Dissection of the flight muscles followed a published protocol ^55^. Briefly, at 48 h APF *Mef2-GAL4, UAS-histone-RFP* pupae were fixed for 30 min in PBS + 0.5% Triton + 4% PFA on a nutator at RT after freeing from their pupal case and piercing 2 holes in the abdomen with dissecting needles. The fixed pupae were pinned on the back with insect needles at the bottom of a silicone Petri dish filled with PBS. A cut was made from the head to the abdomen on both sides to remove the ventral part of the pupa. The tissues of the thorax, except for the flight muscles, were removed with forceps. The two halves of the thorax were cut and transferred to an hourglass. Muscles were labelled for actin by incubation in phalloidin-Alexa488 (1/500) overnight at 4°C and then washed once rapidly with PBST and then 2x in PBST (10 min each) at RT. Muscles were mounted in Vectashield with 1 coverslip spacer. A thick stack (z-step size: 1.16 µm) was imaged to reconstruct the volume of all 6 DLMs from each 48 h APF hemithorax.

#### Data analysis of flight muscles

*Adults:* the actin channel was used in the analysis to delineate the contours of the flight muscles. As adult muscles are too thick to image the entire muscle volume, the volume was estimated by multiplying flight muscle surface area, determined with longitudinal section and muscle thickness determined with transversal section. The number of nuclei was counted using the histone-RFP channel in a large volume of DLMs (250 µm x 250 µm x 40 µm). The z-stack was projected by averaging intensity to visualize all nuclei on a single image. All nuclei present on the projected images were counted and positionally registered. The signal intensity was used to determine the number of nuclei in case of overlap. Since nuclei are uniformly distributed throughout the muscles, the nuclei quantification for the analyzed volume was used to calculate the total number of nuclei for all 12 DLMs: [total DLMs volume] / [analyzed volume] * [number of nuclei from the analyzed volume]. The nuclei density was calculated by dividing the number of nuclei by the volume of the muscle.

*Pupae:* the volume of the developing DLMs at 48 h APF was determined by multiplying the surface area of the muscle on the longitudinal section by the depth of the muscle visualized in the virtual cross-section. The number of nuclei was counted using the histone-RFP channel in a large volume (150 µm x150 µm x 40 µm). Nuclei quantification of 48 h APF DLMs was performed by multiplying the number of nuclei quantified in one longitudinal row with the number of rows of nuclei determined from the 3D volume of the large z-stack. The total number of nuclei was calculated as above for adults. Statistical tests show unpaired t-tests with p-values and each thorax is indicated in the plots.

### Experimental model: organism/strain

*D. melanogaster: Mef2-GAL4* BDSC RRID: BDSC_27390

*D. melanogaster: UAS-histone-mRFP-36* (gift from Frederik Wirtz-Peitz)

### Chemicals

phalloidin-Alexa488 Thermo Fisher Scientific Ref# A12379 Vectashield SKU: H-1000-10

### Experimental validation in heart

#### Fly heart movies

R94C02::tdTomato (attP2) flies were prepared according to the method described ^56^ and illuminated with green light at a power of 3 mW. Five-second recordings of the beating heart were captured at 270 frames per second, and the high-speed movies were analyzed using a custom R script (https://doi.org/10.5281/zenodo.4749935). Fractional shortening (contractility) was calculated as FS = (EDD – ESD) * 100 / EDD, measured from heart chamber A2/A3.

#### Imaging/staining

Adult *w*^1118^ fly hearts were dissected per ^58^ and hearts were stained according to ^59^. In brief, five day old adult female and male flies were anaesthetized and placed on their dorsal side on a dish covered with a layer of petroleum jelly. After addition of artificial hemolymph, flies were dissected to expose the heart tube, followed by formaldehyde fixation and permeabilization with PBTx. The hearts were stained with antibodies for H15/Neuromancer-1 ^60^ [guineapig, 1:2000], Fibrillarin [Mouse, 38F3 from EnCor Biotechnology, 1:100] and for F-actin (phalloidin, 1:1000, Invitrogen) and mounted in DAPI-containing Prolong Gold antifade (Invitrogen). Adult heart stacks were acquired on a Zeiss Imager.Z1 equipped with an Apotome2 and Hamamatsu Orca Flash4 using a LCI Plan-Neofluar 25x/0.8 objective and 1.8µm Z-step size.

#### Nuclear feature annotation and data analysis

Heart stacks were imported as native Zeiss CZI files into napari v. 0.4.17 ^61^ and the cardiac nuclei channel labeled by H15, all nuclei (DAPI) and all nucleoli (Fibrillarin) were segmented using the Accelerated Pixel and Object Classification (APOC) plugin v.0.12.3 ^62^. Pericardial cells (PCs) can be morphologically identified and were manually annotated. Label coordinates were exported as CSV files and processed in R using a custom script (https://zenodo.org/doi/10.5281/zenodo.13820362) to match cardiac and PC nuclei with their respective nucleoli and to calculate cell type specific feature volumes and ratios.

### Experimental validation in fat body

#### Fly Husbandry

Larvae were raised at a density of 50 animals per vial at 25°C in *Drosophila* growth medium consisting of 20.5g/L sucrose, 70.9g/L D-glucose, 48.5 g/L cornmeal, 60.6 g/L yeast, 4.55 g/L agar, 0.5 g CaCl2.2H2O, 0.5 g MgSO4.7H2O, 11.77 mL acid mix (propionic acid/phosphoric acid). Adult flies were sexed and raised separately at a density of 20 flies per vial. For all experiments, equal numbers of similarly sized and developmentally staged larvae or adults were used per experimental group.

#### Immunostaining and microscopy

Larvae were sexed according to gonad size and dissected 108 hours after egg laying. Inverted larvae were fixed in 4% paraformaldehyde in phosphate-buffered saline (PBS) for 1 hr. Carcasses were rinsed once in PBS for 10 mins and thrice in 0.1% Triton-X in PBS (PBST) for 5 mins. Carcasses were then blocked in a solution of 3% Normal Goat Serum (NGS) in 0.1% PBST for 1 hour at room temperature and incubated with the monoclonal antibody to fibrillarin at a dilution of 1:400 overnight at 4°C. The carcasses were washed with 0.1% PBST three times for 5 mins each and then incubated with goat anti-rabbit secondary antibody conjugated to Alexa-Fluor 568 or 488 (1:1000) for 2 hrs. Carcasses were washed thrice with 0.1% PBST again where the penultimate wash was extended to stain with Hoechst 3342 before mounting in SlowFade Diamond mounting medium. For quantification of nuclear and nucleolar area, images were captured with a Leica SP5 laser scanning confocal microscope.

#### Protein synthesis assay

Larvae were inverted in *Drosophila* Schneider’s medium and then incubated with 20 uM OPP reagent solution for 30 mins. Carcasses were fixed in 4% paraformaldehyde in 1X phosphate-buffered saline (PBS) for 1 hour and then rinsed thrice in PBS for 5 mins each. The rest of the assay was performed using the Click-iT Plus OPP Alexa Fluor 488 kit (Invitrogen; C10456) according to manufacturer’s instructions. The larval fat bodies were dissected out of the carcasses and mounted in SlowFade Diamond mounting medium. Images were acquired with a Leica SP5 laser scanning confocal microscope and Fiji (version 1.53q) was used for quantification of OPP staining intensity. One biological replicate represents the average of total fluorescent intensity per cell of 5 cells in a fat body. All intensity measurements were normalized to cell volume. 15 different larvae per sex were measured.

#### RNA isolation and cDNA synthesis

For the larval fat body, each biological replicate represents 15 fat bodies from 15 larvae. For the adult fat body, 15 fly abdominal carcasses dissected at 5 days post-eclosion make one biological replicate. The fat bodies were collected into 1 ml of TRIzol (Thermo Fisher Scientific; 15596018) in 1.5-ml microcentrifuge tubes. All samples were frozen on dry ice and stored at −80°C until processing. Each experiment contained seven biological replicates per sex, and each experiment was performed in two sets with four replicates in the first set and three replicates in the second. Total RNA was extracted as previously described ^3,63–65^. Genomic DNA was eliminated, and cDNA was synthesized using the QuantiTect Reverse Transcription Kit (Qiagen) according to the manufacturer’s instructions.

#### qPCR

qPCR was performed in a 15-μL reaction volume containing 2 μL of cDNA and final concentrations of 0.3 U of recombinant Taq DNA Polymerase (Thermo Fisher Scientific), 0.1× SYBR Green I Nucleic Acid Gel Stain (Thermo Fisher Scientific), 0.3 μM of specific primer pairs (Integrated DNA Technologies, Eurofin Genomics, Thermo Fisher Scientific), 1× PCR buffer (Thermo Fisher Scientific), 125 μM dNTP mix (FroggaBio), and 1.5 mM MgCl2 (Thermo Fisher Scientific). qPCR was carried out in a CFX384 Touch Real-Time PCR Detection System (BioRad). Thermocycler conditions were as follows: initial denaturation for 3 minutes at 95°C and then 40 cycles of denaturation for 30 seconds at 95°C, annealing for 30 seconds at 60°C, and extension for 45 seconds at 72°C.

To show the sex bias in gene expression, Cq values were normalized to the average of female Cq values (ΔCq) and fold change was calculated as 2(absolute value of ΔCq). Data were then normalized to the average fold change of actin and β-tubulin.

#### Larval fat cell quantification and body size

Pupal volume was measured in male and female pupae as previously described ^3,63,64,66^. Pupae were tracked individually until they eclosed.

Larval fat cells were quantified as previously described ^65^. Briefly, newly eclosed single male or female fly was dissected open through the abdomen in one well of a 24-well plate using 500 μl of 1X PBS with 1:100 dilution of BODIPY 493/503 (Thermo Fisher, D3922) and 1:500 dilution of Hoechst 33342 (Thermo Fisher, H3570) in 1X PBS. The carcass was removed once the larval fat cells were loosened from the carcass into the well. After 40 min of incubation at room temperature, fat cells images were acquired using Zeiss AXIO ZoomV.16. ImageJ was used to count the number of larval fat cells within each well. The number of larval fat cells for every individual fly was normalized to its pupal volume.

### Visualization

The analysis and visualization were performed using a combination of R v4.2.2 ^67^ and Python v3.11.10, with all scripts available in the Zenodo repository https://zenodo.org/doi/10.5281/zenodo.13820362. Scatter plots for count bias were generated using the `ggplot2` v3.4.1 package from the `tidyverse` v2.0.0 collection in R. Heatmaps for RP and other related genes were created using the `ComplexHeatmap` v2.15.4 package.

Experimental boxplots were produced using the ggplot2 package, with significance bars added via the ggsignif v0.6.4 package, utilizing the geom_signif function and setting the test parameter to t.test.

## Acknowledgements

We thank the Przytycka and Oliver Lab members for their contributions during lab meeting discussions.

## Author contributions

Each author’s contribution(s), according to the CRediT model, is listed below.

Conceptualization: SP, TMP, BO

Methodology: SP, EJR, FS, GV, TMP, BO

Investigation: SP, JA, CMC, PB, GV

Visualization: SP, JA, CMC, EJR, GV, BO

Funding acquisition: EJR, FS, TMP, BO

Project administration: SP, EJR, FS, TMP, BO

Supervision: EJR, FS, TMP, BO

Writing – original draft: SP, JA, CMC, EJR, FS, GV, TMP, BO

Writing – review & editing: SP, JA, CMC, EJR, FS, GV, TMP, BO

## Data and materials availability

The single-cell Fly Cell Atlas data analyzed in this study is available at https://www.flycellatlas.org. All code used for analyzing FCA data, along with the experimental data and related analyses, as discussed in this paper, are accessible via the Zenodo repository: https://zenodo.org/doi/10.5281/zenodo.13820362.

## Supplementary Materials

Supplementary Text

Figs. S1 to S19

Tables S1 to S7

Data S1 to S14

