## Supplementary material for "Cell type specific allometry controls sex-differences in *Drosophila* body size": Suppl Text

**The PDF file includes:**

Figs. S1 to S19
Tables S1 to S7

**Other Supplementary Materials for this manuscript include the following:**

Data S1 to S14

**Fig. S1.**

Scatter plots of the FCA Leiden 1.0 clusters for dissected tissues: (**A**) Antenna, (**B**) Body Wall, (**C**) Fatbody, (**D**) Gut, (**E**) Haltere and (**F**) Heart. The Y-axis represents the percentage of female cells, and the X-axis represents the percentage of male cells, both normalized by the total cell count in the replicated samples. The dashed line indicates equal proportions of female and male cells, while the dotted lines show a cell proportion bias of 1.25-fold, and the solid lines represent a bias of 2.0-fold. Female-biased clusters are shown in red, and male-biased clusters in blue, with darker shades representing a bias of at least 2.0-fold. Clusters containing more than 1% of cells are color-coded based on their nine broad annotations, while those below this threshold are shown in gray. Only a few selected clusters are shown with their major annotation.

**Fig. S2.**

Scatter plots of the FCA Leiden 1.0 clusters for dissected tissues: (**A**) Leg, (**B**) Malpighian Tubule, (**C**) Oenocyte, (**D**) Proboscis and Maxillary Palps, (**E**) Trachea and (**F**) Wing. The Y-axis represents the percentage of female cells, and the X-axis represents the percentage of male cells, both normalized by the total cell count in the replicated samples. The dashed line indicates equal proportions of female and male cells, while the dotted lines show a cell proportion bias of 1.25-fold, and the solid lines represent a bias of 2.0-fold. Female-biased clusters are shown in red, and male-biased clusters in blue, with darker shades representing a bias of at least 2.0-fold. Clusters containing more than 1% of cells are color-coded based on their nine broad annotations, while those below this threshold are shown in gray. Only a few selected clusters are shown with their major annotation.

**Fig. S3-S16.**

Heatmaps showing the expression bias of ribosomal protein genes (RP; FBgg000141) and a small set of transcription-related genes grouped as eEF, eIF, eRF, mitochondrial, and special categories for the head (S3) and body (S4) datasets, as well as 12 dissected tissue datasets (S5-16). Genes are shown as rows and Leiden clusters as columns. All heatmaps use a consistent color scale.

**Fig. S17.**

Sex-specific expression bias in ribosomal proteins (as a proxy for cell size) and additional genes related to translation. (**A-B**) Heatmaps displaying pairwise Pearson's correlation of expression bias for all ribosomal protein genes (FBgg0000141) show a strong correlation in sex-biased expression among ribosomal protein-encoding genes within cell types in the head (**A**) and headless body (**B**) samples. This suggests a coordinated effort by genes in the ribosome, a large multi-subunit complex (Fig. 3A), to produce more ribosomes in females. (**C-D**) Several eukaryotic initiation factors (eIFs; Table S2) and eukaryotic elongation factors (eEFs; Table S2) exhibit significantly higher female-biased expression in both the head (**C**) and headless body (**D**) samples. Both heatmaps use a consistent color scale. At the bottom of both heatmaps, we show the expression bias of additional translation-related genes: DENR, MCTS1, Nelf-A, NELF-B, Nelf-E, eRFs, pix, and tor. (**E**) Quantitative real-time PCR (qPCR) revealed elevated relative mRNA levels for 14 ribosomal protein genes in the larval fat body, similar to Fig. 3F for the adult fat body. Each box plot displays the first and third quartiles as the hinges, with the median represented by the line in the middle. The upper whisker extends from the upper hinge to the largest value within 1.5 times the interquartile range (IQR) from the hinge (IQR is the distance between the first and third quartiles). Similarly, the lower whisker extends from the lower hinge to the smallest value within 1.5 times the IQR. p-values are calculated using the t-test.

**Fig. S18. Extended figures on sexual allometry of the heart.**

(**A**) Sex differences are reflected in end-systolic diameter (ESD) similar to end-diastolic diameter (EDD; Fig. 3A-B). (B) Sex bias in cell counts in heart FCA clusters. Note the clusters are as annotated in FCA unlike Fig. S1F where clusters are at Leiden 1.0 resolution (C) Sex bias in UMI counts in cardiomyocytes in the FCA heart sample shows a female bias in ribosomal protein genes. Ribosomal protein genes are shown in blue, whereas other genes are shown in gray. The red dotted line represents equal counts.

**Fig. S19. Extended figures on sexual allometry of the fat body.** (**A**) Representative images of $w^{1118}$ female (top row) and male (bottom row) larval fat body dissected at 108 hours after egg-laying (AEL), showing fibrillarin-positive nucleoli (left column), Hoechst-stained nuclei (middle column), and the merged images (right column). Scale bars: 20 µm.

**Table S1.**

Excel workbook providing details about the single-cell Fly Cell Atlas (FCA) samples and annotated cell types used in this study. It contains three spreadsheets: (1) **ReadMe**: Describes the columns in the other two sheets. (2) **FCA_samples**: Lists details about FCA loom file URLs and the resolutions used in this study. (3) **FCA_Annotations**: Includes the list of annotated cell types, shorter annotations, a broad category of nine cell types the annotation belongs to and an indication whether they were classified as female-specific, male-specific, or non-sex-specific. For this study, we used only non-sex-specific cells and excluded those annotated as artefacts.

**Table S2.**

Excel workbook providing details about the ribosomal and other translation-related genes. It contains two spreadsheets: (1) **ReadMe**: Explains the columns in the other sheet. (2) **SymbolsGroups**: Lists the gene symbols and their groupings as presented in the heatmaps.

**Table S3.**

Excel workbook providing details on cluster sex bias in both cell counts and gene expression for the genes listed in Table S2. It contains fifteen spreadsheets: (1) **ReadMe**: Explains the columns in the other fourteen sheets. (2-15): Each corresponds to one of the FCA samples listed in Table S1, with cluster information in the columns and gene information (from Table S2) in the rows.

**Table S4.**

Consolidated list and details of the reagents and software utilized in this study (see Materials and Methods section).

**Table S5.**

Excel workbook providing details about the experimental data from indirect flight muscle. It contains seven spreadsheets: (1) **ReadMe**: Explains the columns in the other sheet. (2) **MuscleSizeAdult**: Volume of nucleus of adult indirect flight muscle, (3) **MuscleSize48hAPF**: Volume of nucleus of larval (48h APF) indirect flight muscle, (4) **MuscleNucleiCountAdult**: Number of nucleus of adult indirect flight muscle, (5) **MuscleNucleiCount48hAPF**: Number of nucleus of larval (48h APF) indirect flight muscle, (6) **MuscleNucleiDensityAdult**: Nuclear density of adult indirect flight muscle, (7) **MuscleNucleiDensity48hAPF**: Nuclear density of larval (48h APF) indirect flight muscle.

**Table S6.**

Excel workbook containing detailed experimental data from the heart. It consists of three spreadsheets: (1) **ReadMe**: Describes the columns in the other sheets; (2) **HeartNuclearNucleolarVolume**: Provides estimated volumes of the nucleus and nucleolus in three heart cell types; and (3) **HeartStatistics**: Includes various statistics on fly heart, such as systolic and diastolic diameters.

**Table S7.**

Excel workbook containing detailed experimental data from the fat body. It consists of seven spreadsheets: (1) **ReadMe**: Describes the columns in the other sheets; (2) **FatbodyCellCount**: Volume and number of cells in fat body samples; (3) **FatbodyArea**: Nuclear and nucleolar area of cells in fat body samples; (4) **FatbodyRPgeneAdult**: mRNA levels of large (RpL), small (RpS), and mitochondrial large (mRpL) ribosomal subunit genes in $w^{1118}$ adult abdominal carcasses; (5) **FatbodyRPgeneLarva**: mRNA levels of large (RpL), small (RpS), and mitochondrial large (mRpL) ribosomal subunit genes in $w^{1118}$ larval abdominal carcasses; (6) **FatbodyMycRNAi**: Nuclear and nucleolar area of cells in fat body Myc-RNAi samples; and (7) **FatbodyProteinSynthesis**: Fluorescent intensity of synthesized proteins in fat body samples.

**Data S1-14. (separate files)**

Comprehensive information regarding sex differences in cell counts and gene expression is provided for each of the 14 samples in H5AD format. Please note that Table S4 presents only a portion of this data in Excel format. Complete details about the row and column information can be found in the source code at (https://zenodo.org/doi/10.5281/zenodo.13820362).
