## Supplementary figures and images for "Cell type specific allometry controls sex-differences in *Drosophila* body size"

### Supp Fig S1

#### Figure S1 - Count Bias in first 6 dissected tissues

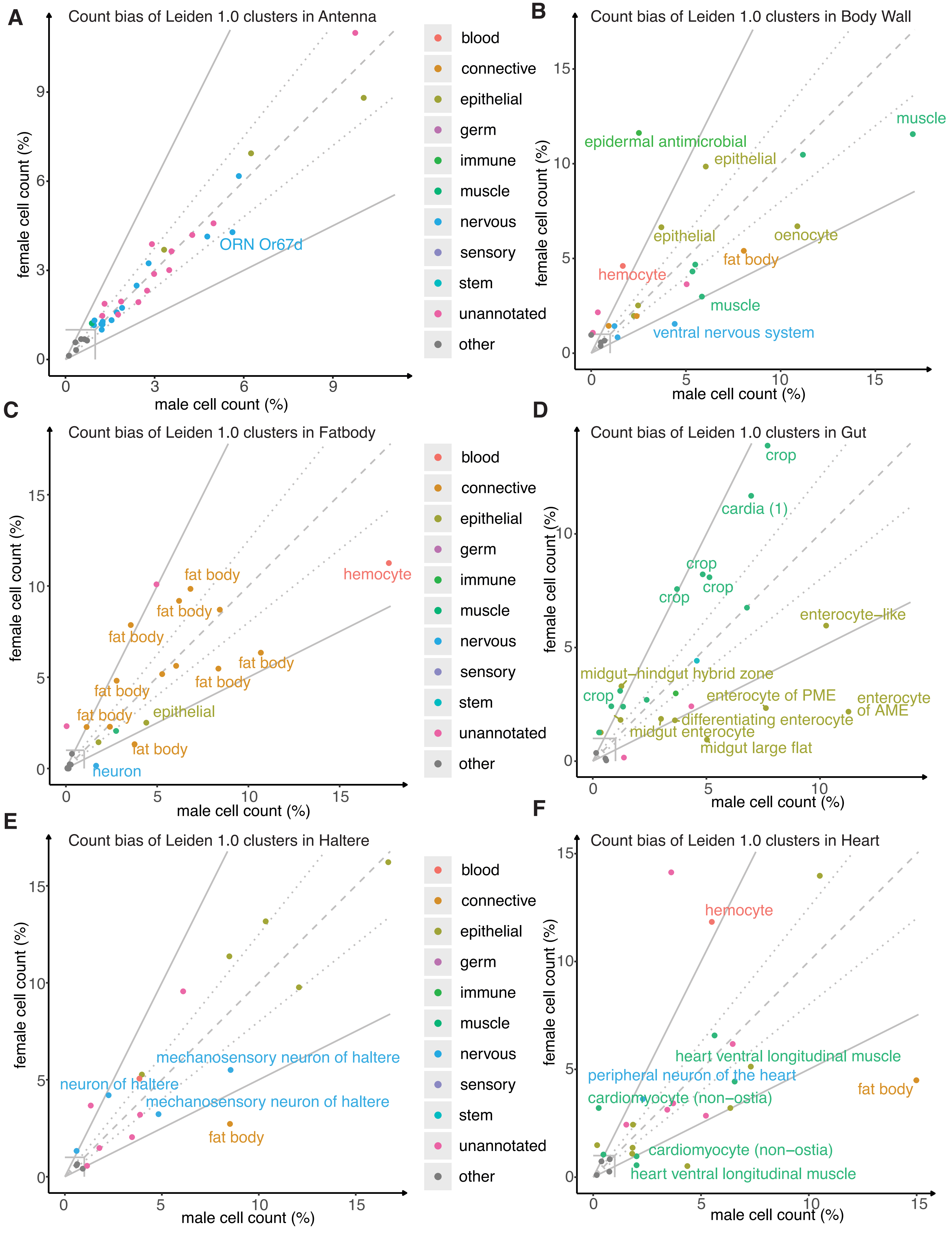

### Supp Fig S2

#### Figure S2 - Count Bias in last 6 dissected tissues

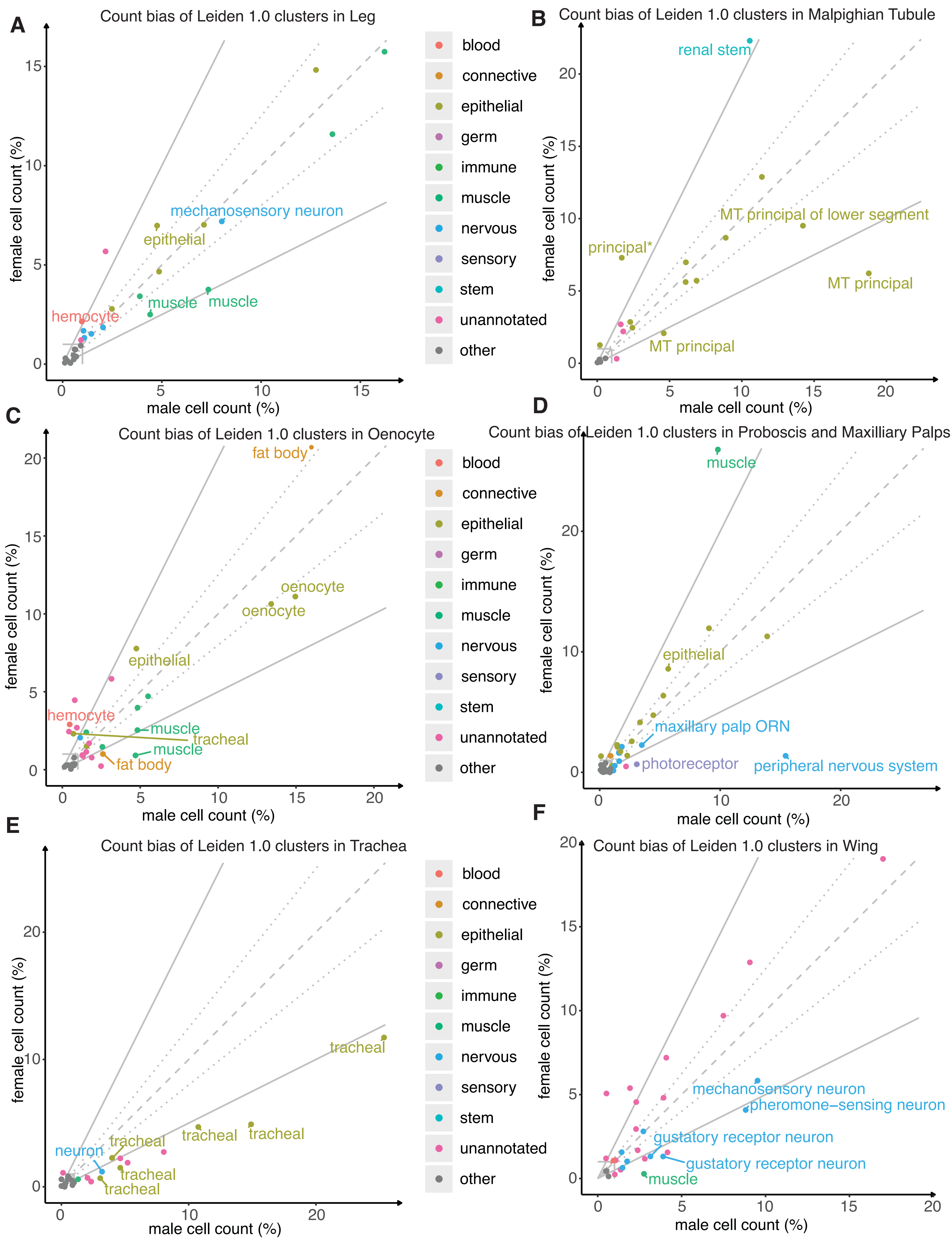

### Supp Fig S3

#### Figure S3 - Expression Bias in Head

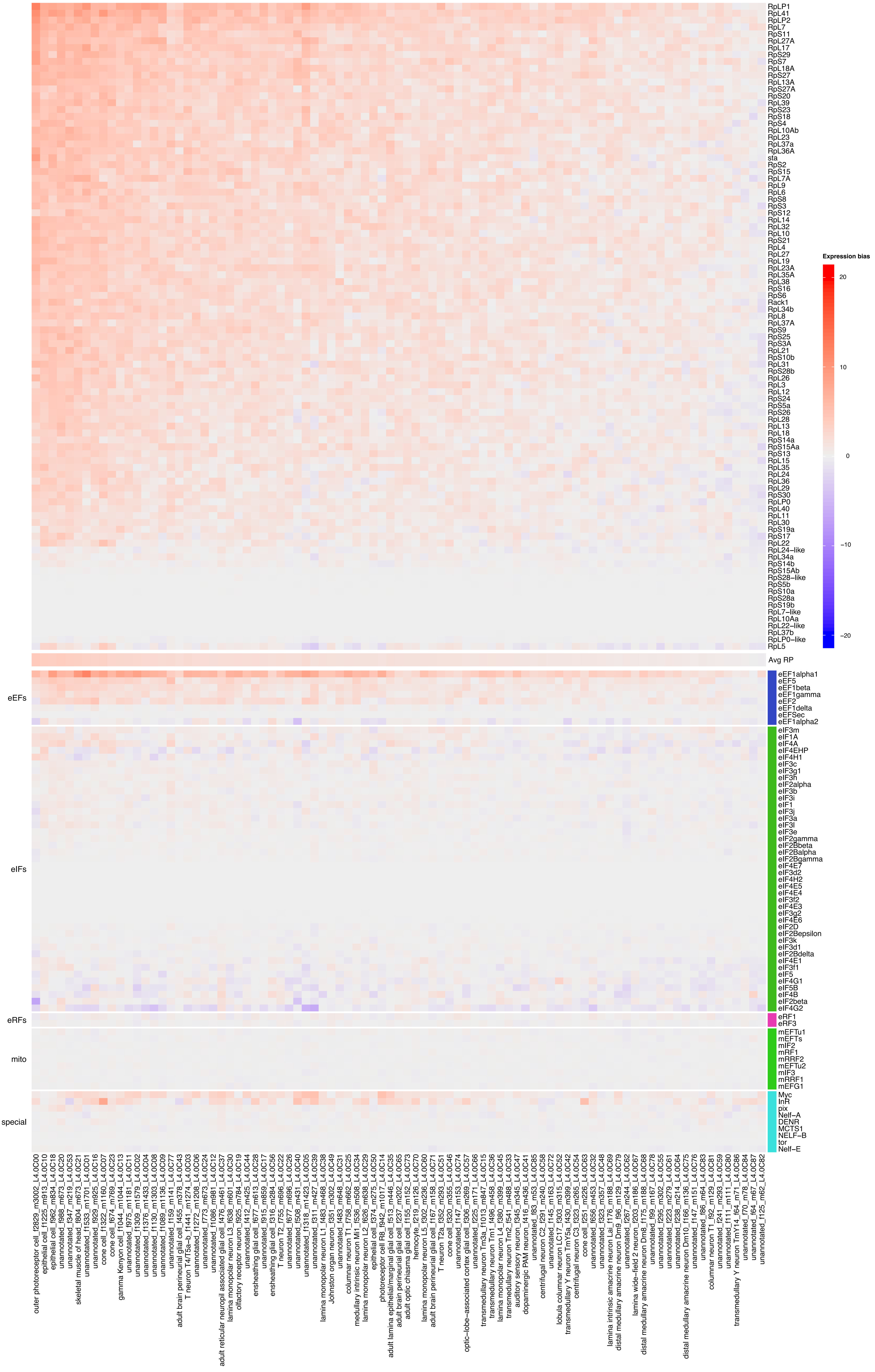

### Supp Fig S4

#### Figure S4 - Expression Bias in Body

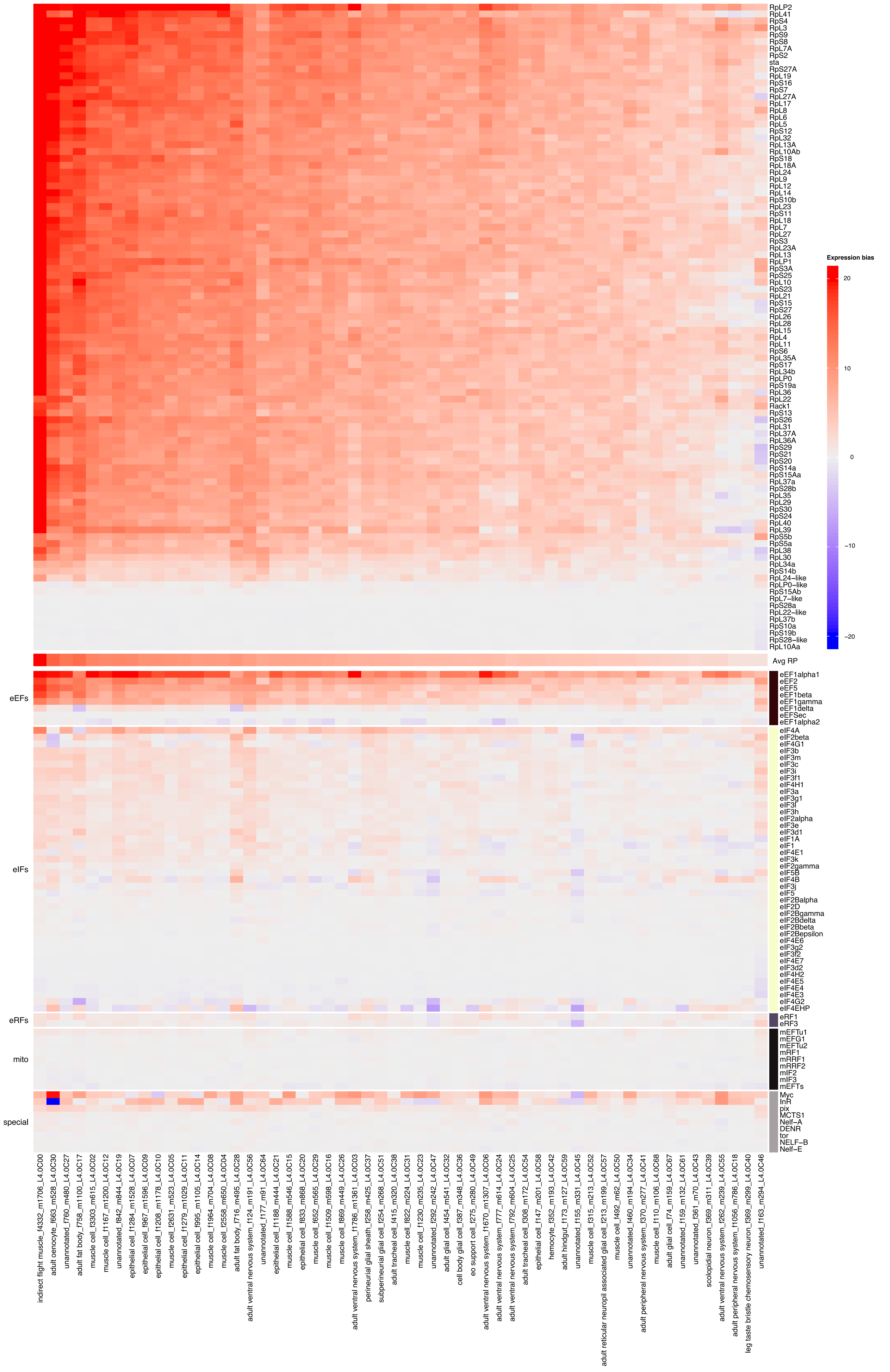

### Supp Fig S5

#### Figure S5 - Expression Bias in Antenna

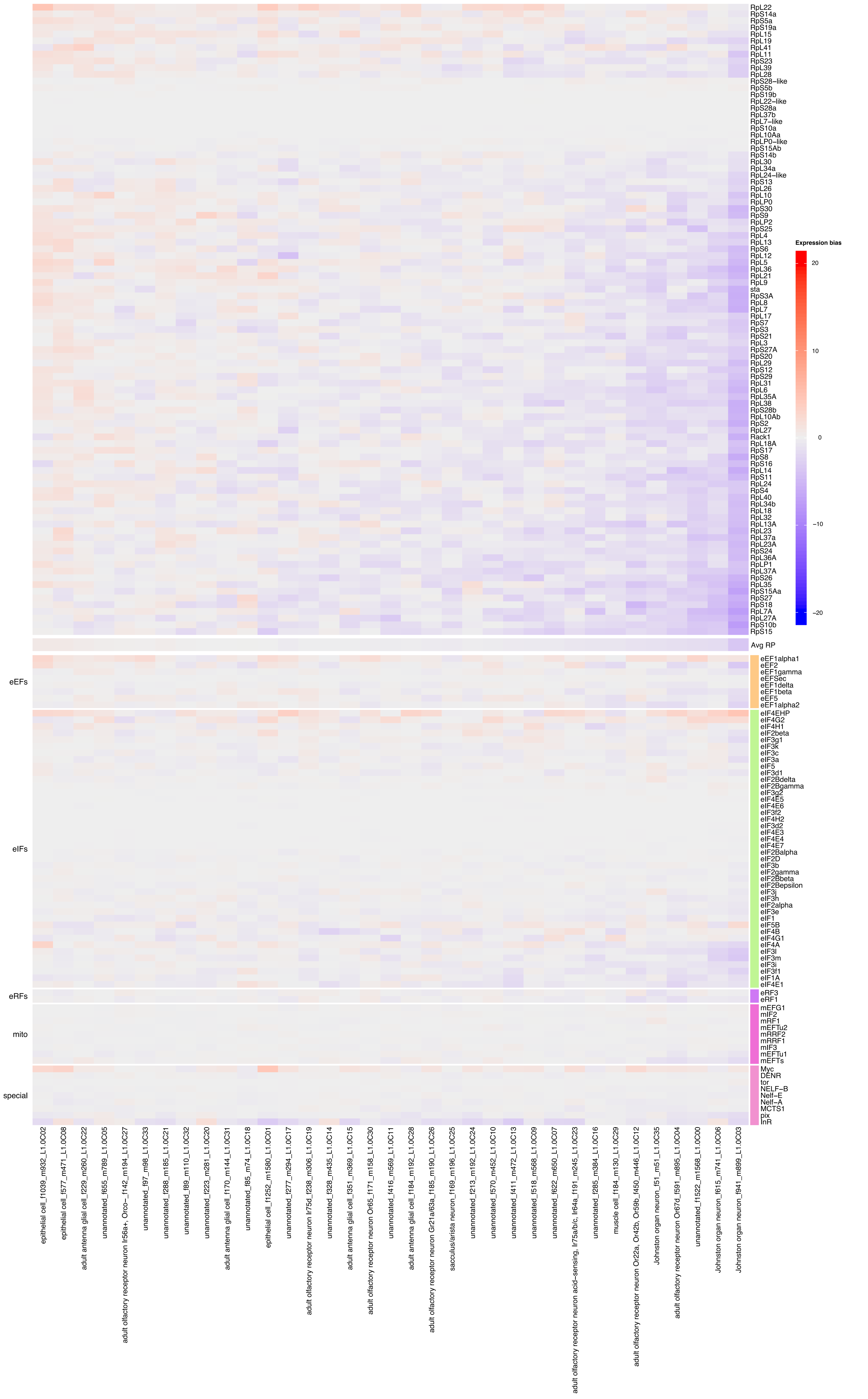

### Supp Fig S6

#### Figure S6 - Expression Bias in Body Wall

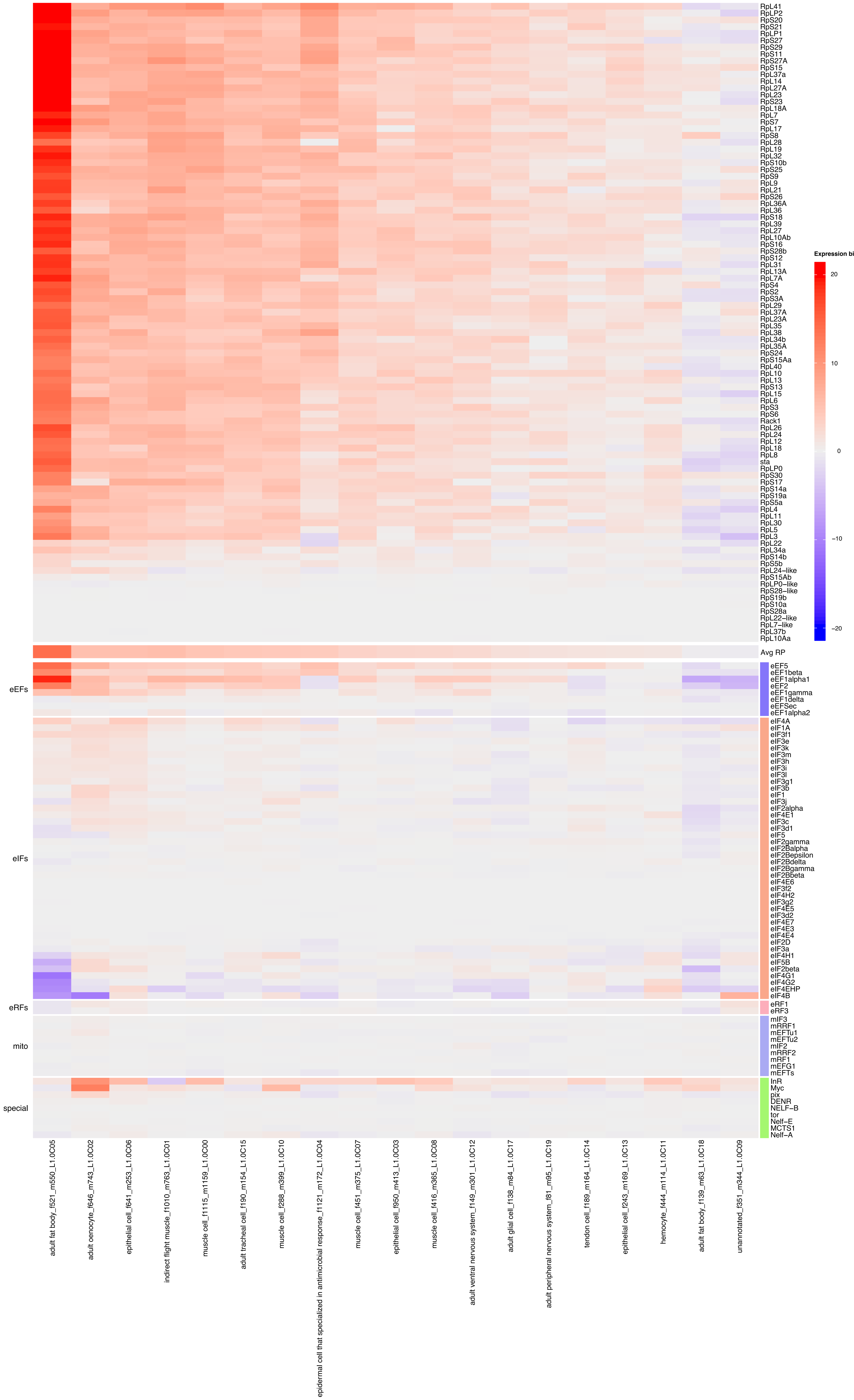

### Supp Fig S7

#### Figure S7 - Expression Bias in Fatbody

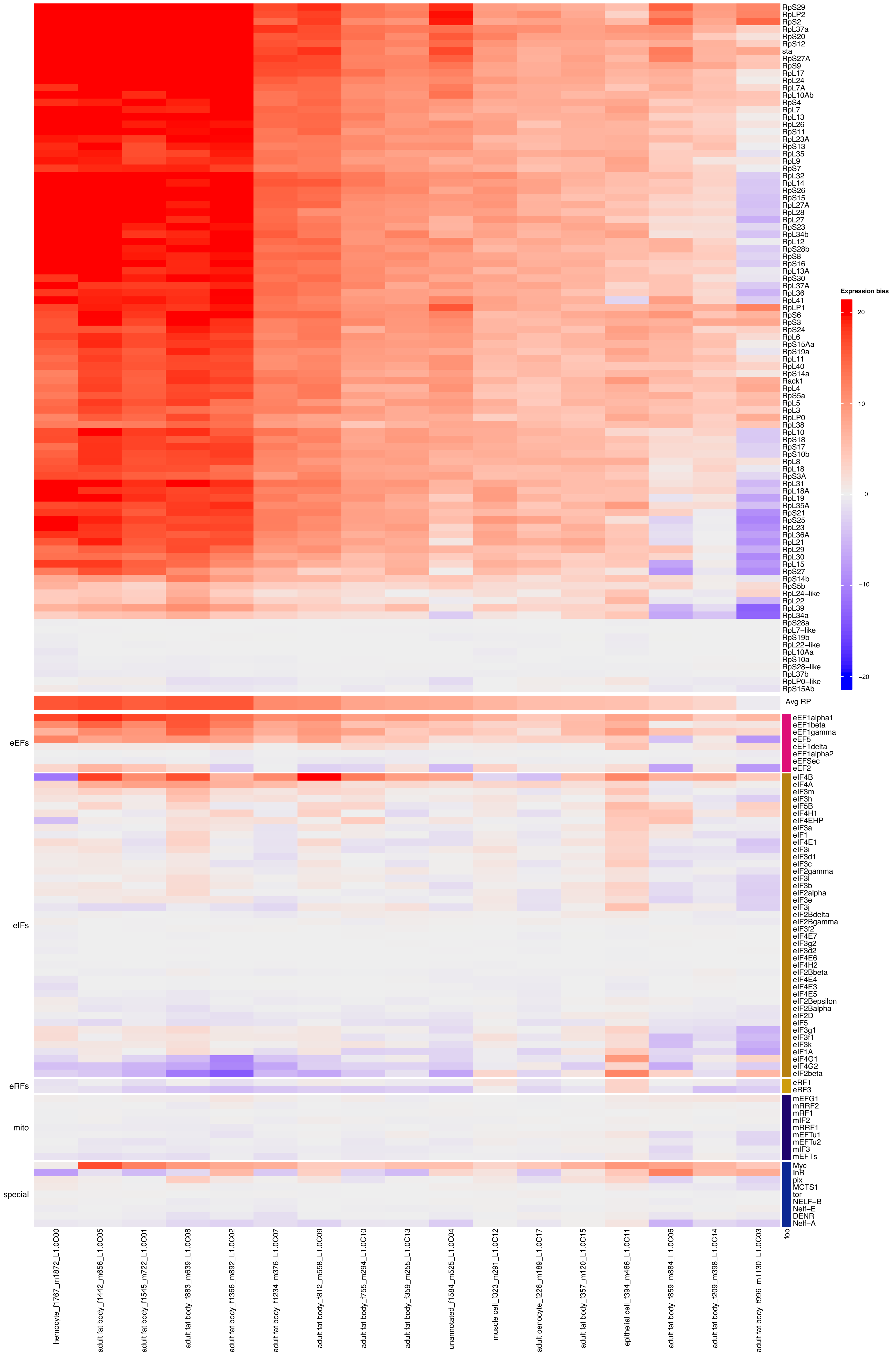

### Supp Fig S8

#### Figure S8 - Expression Bias in Gut

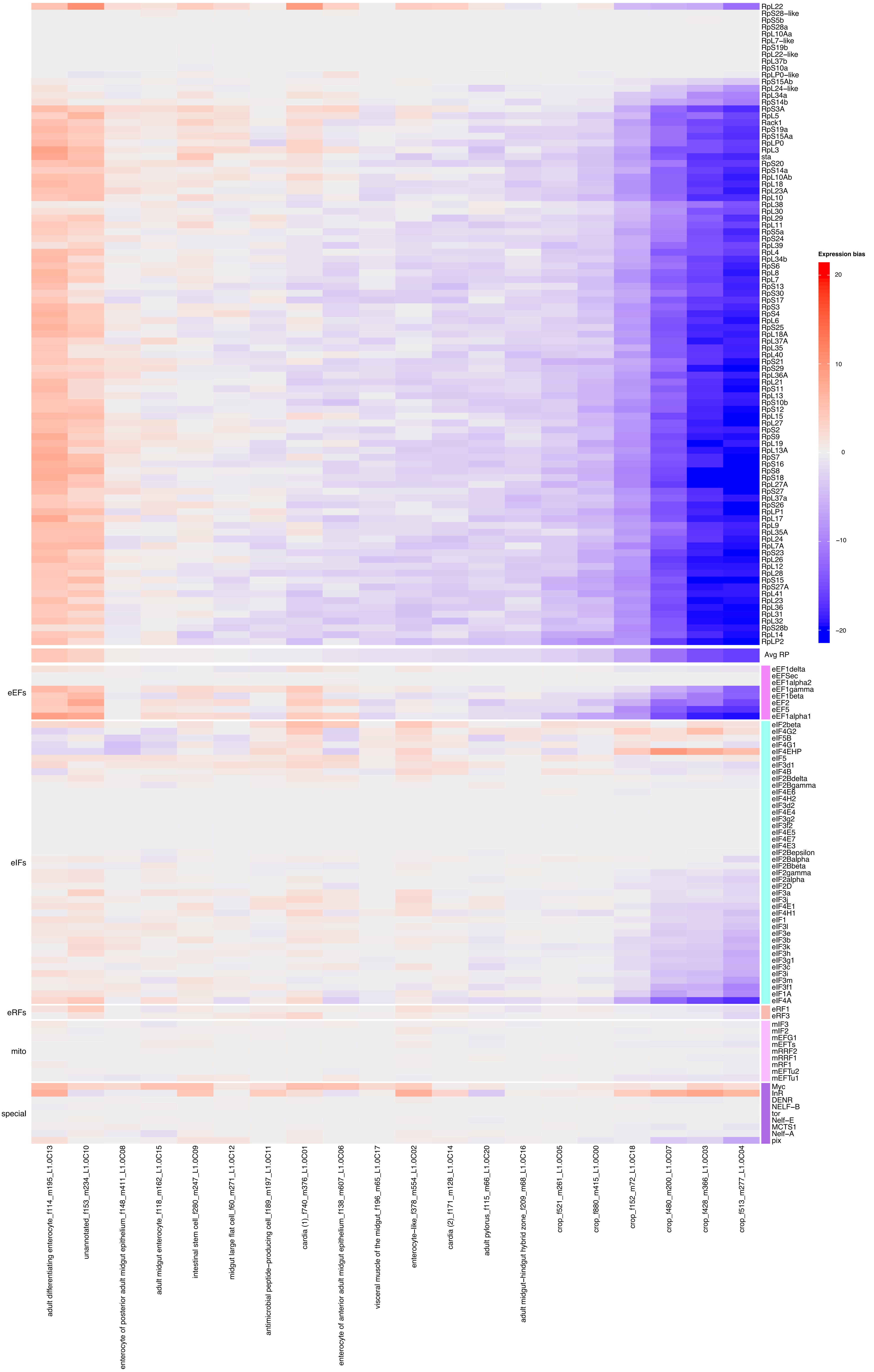

### Supp Fig S9

#### Figure S9 - Expression Bias in Haltere

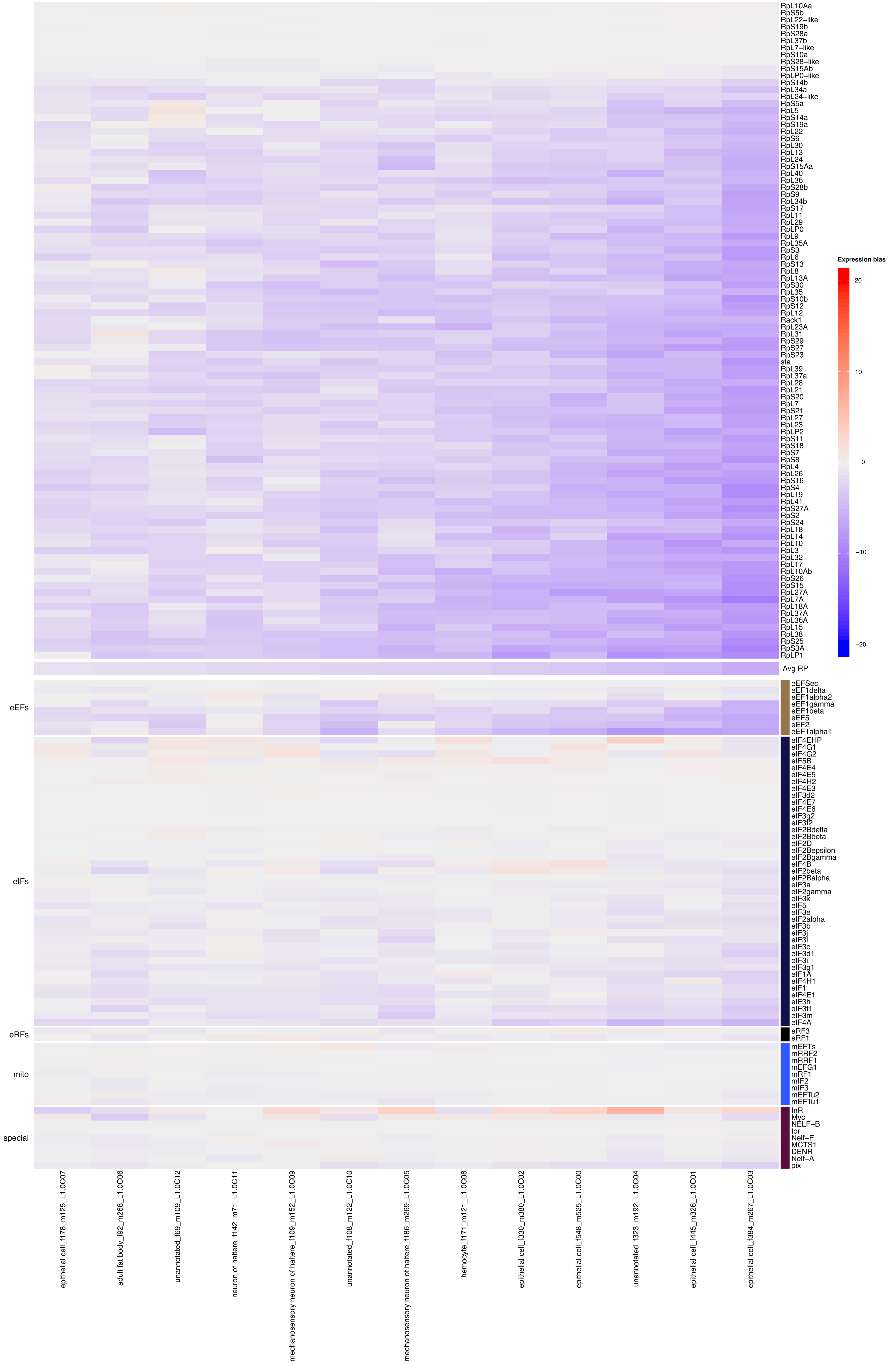

### Supp Fig S10

#### Figure S10 - Expression Bias in Heart

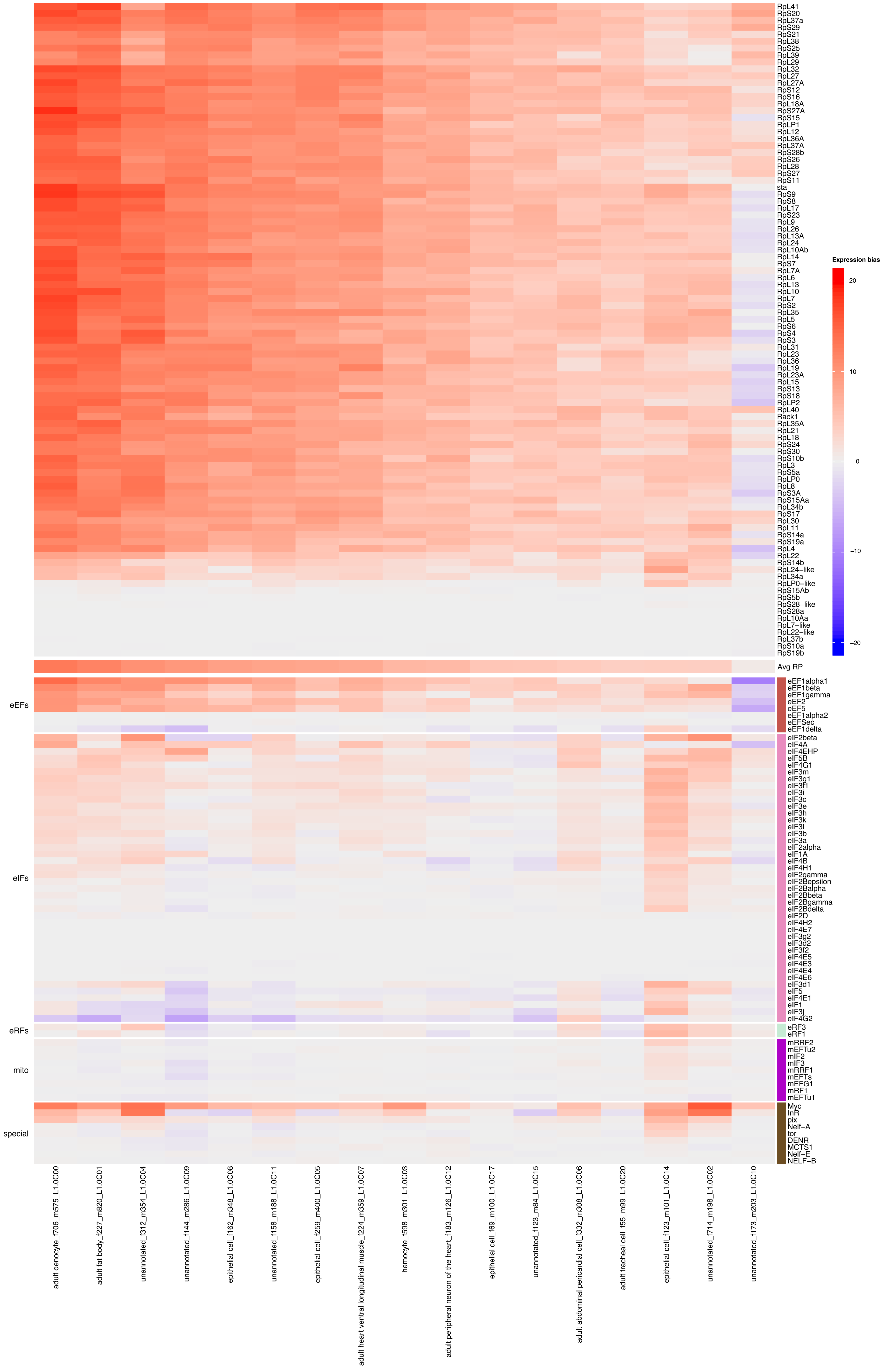

### Supp Fig S11

#### Figure S11 - Expression Bias in Leg

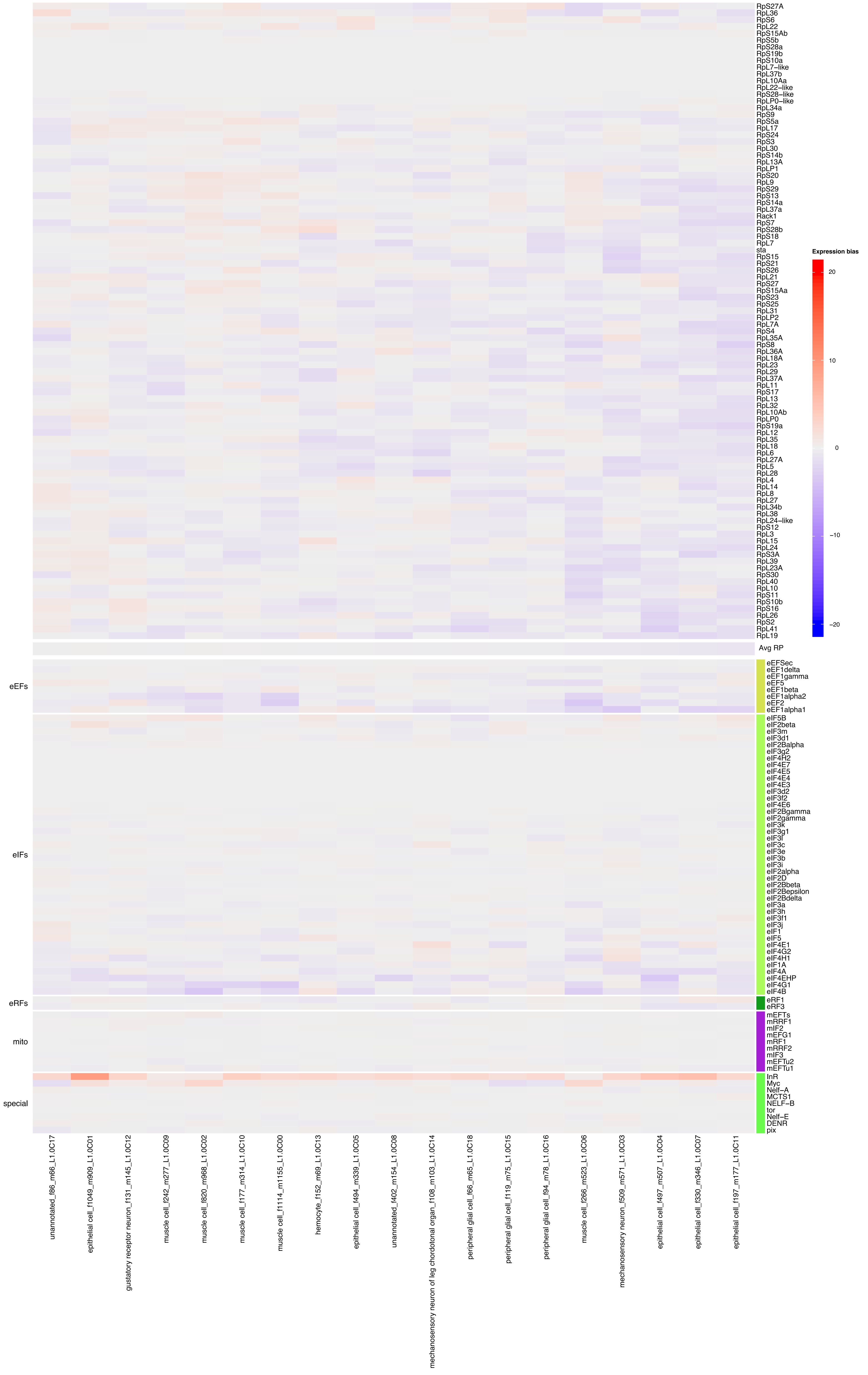

### Supp Fig S12

#### Figure S12 - Expression Bias in Malpighian Tubule

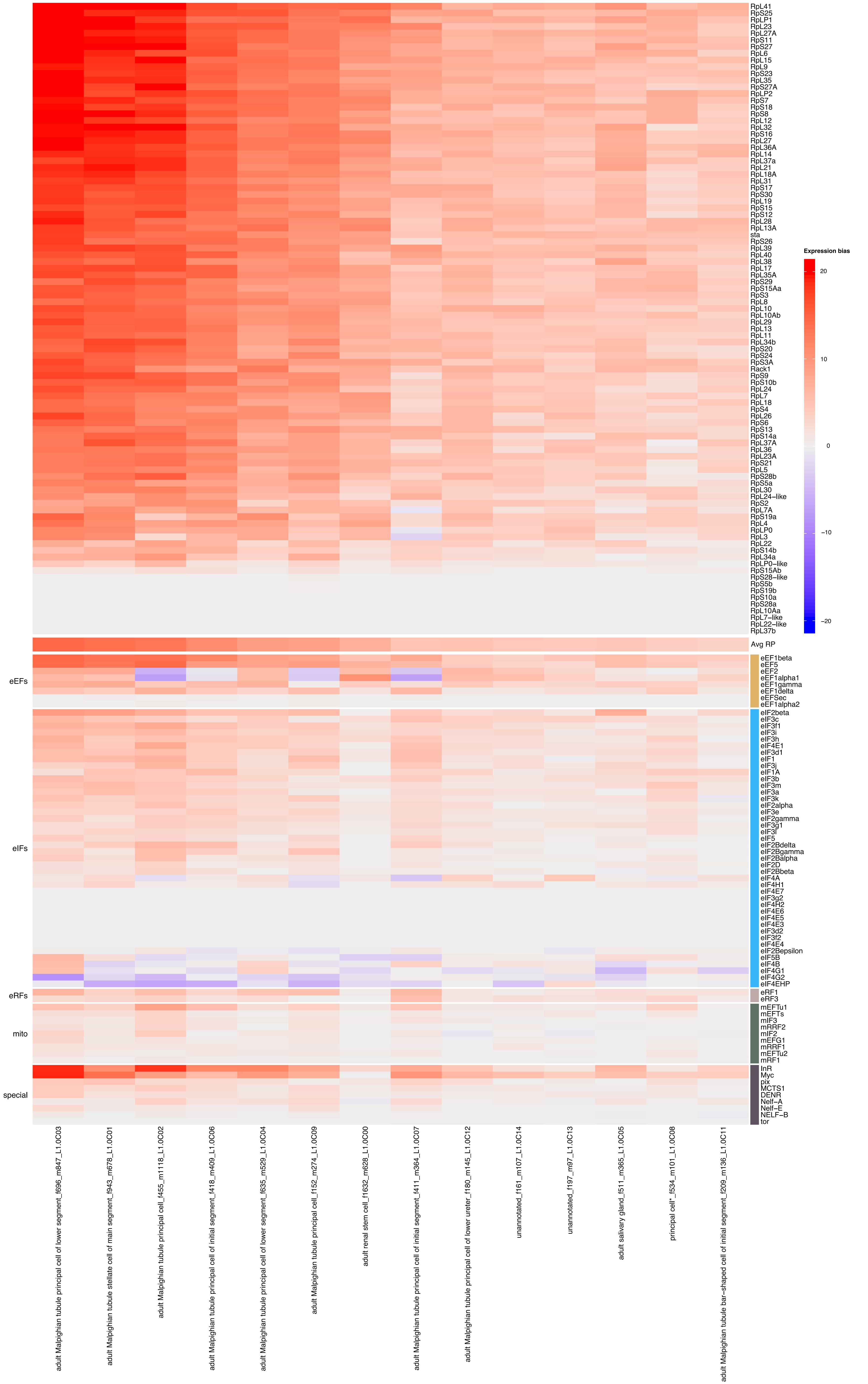

### Supp Fig S13

#### Figure S13 - Expression Bias in Oenocyte

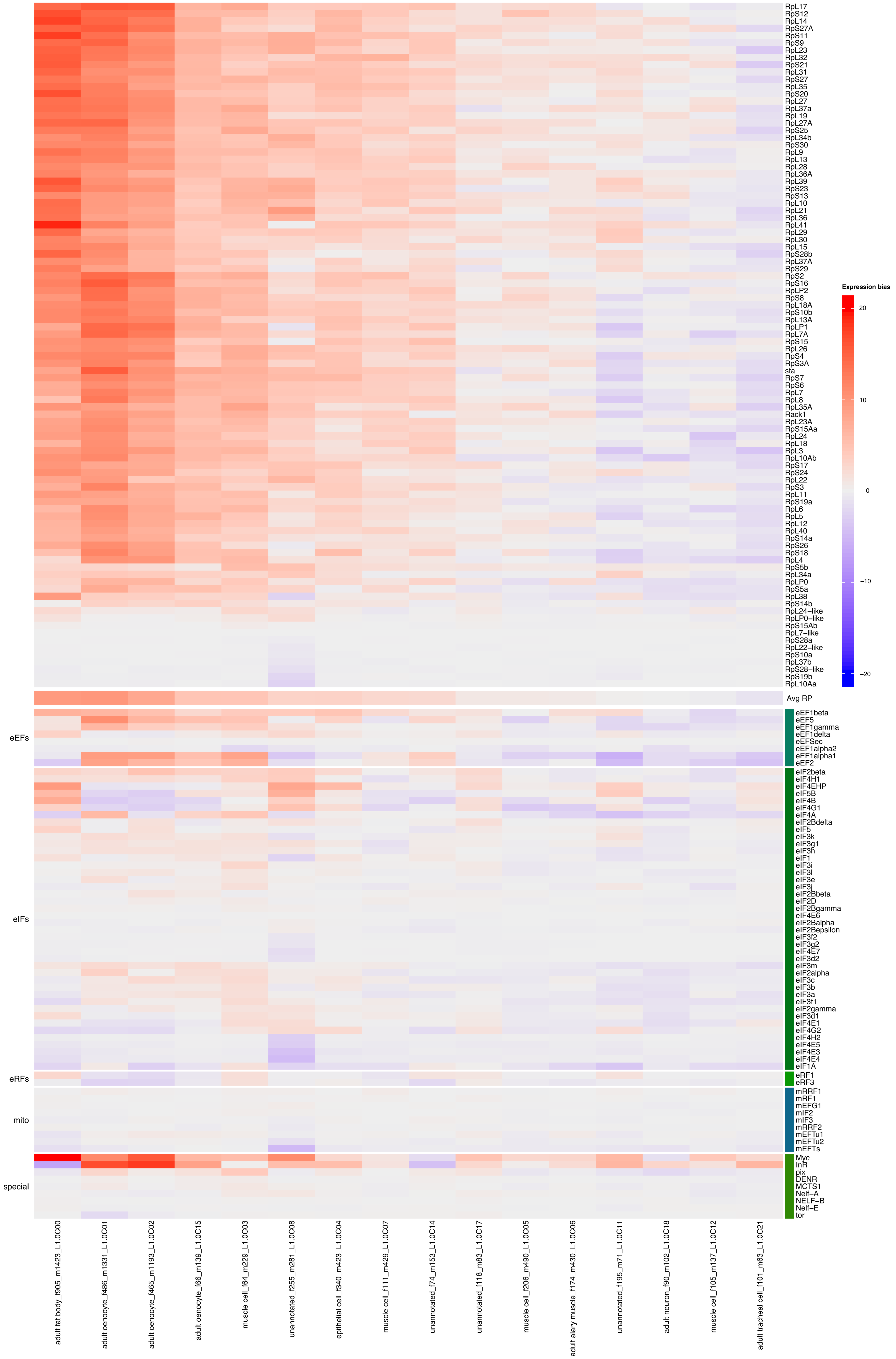

### Supp Fig S14

#### Figure 14 - Expression Bias in Proboscis and Maxillary Palps

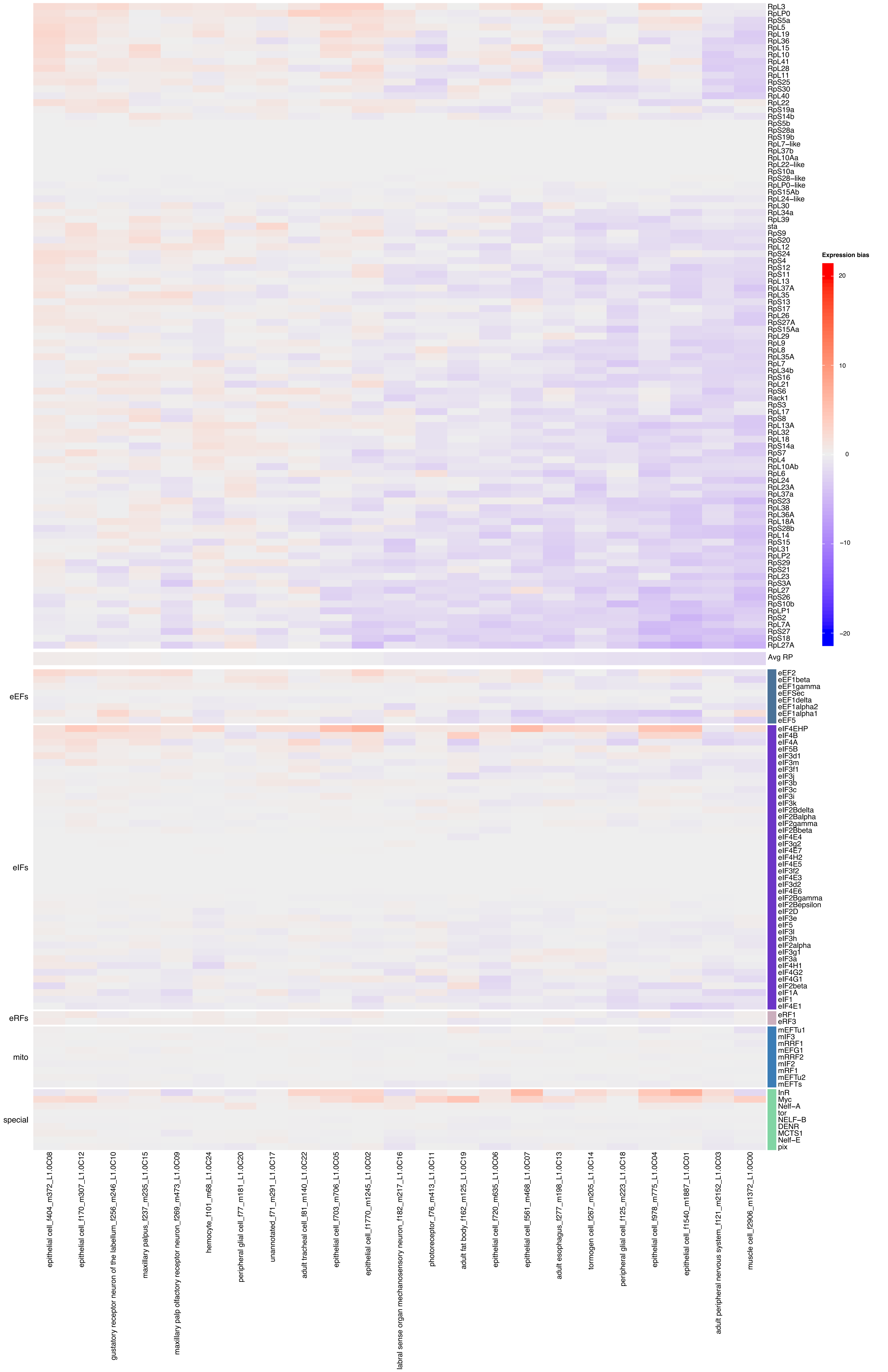

### Supp Fig S15

#### Figure S15 - Expression Bias in Trachea

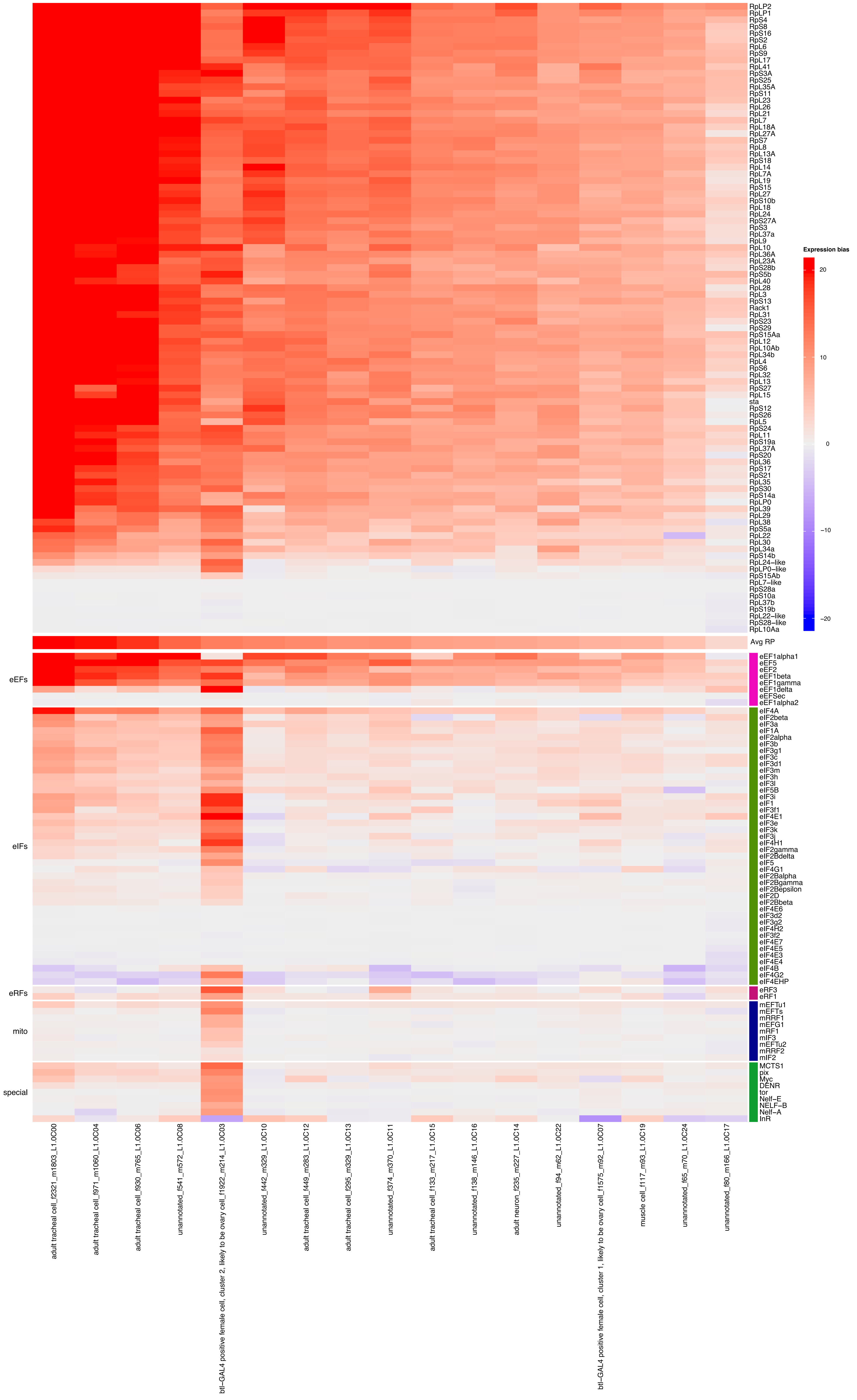

### Supp Fig S16

#### Figure S16 - Expression Bias in Wing

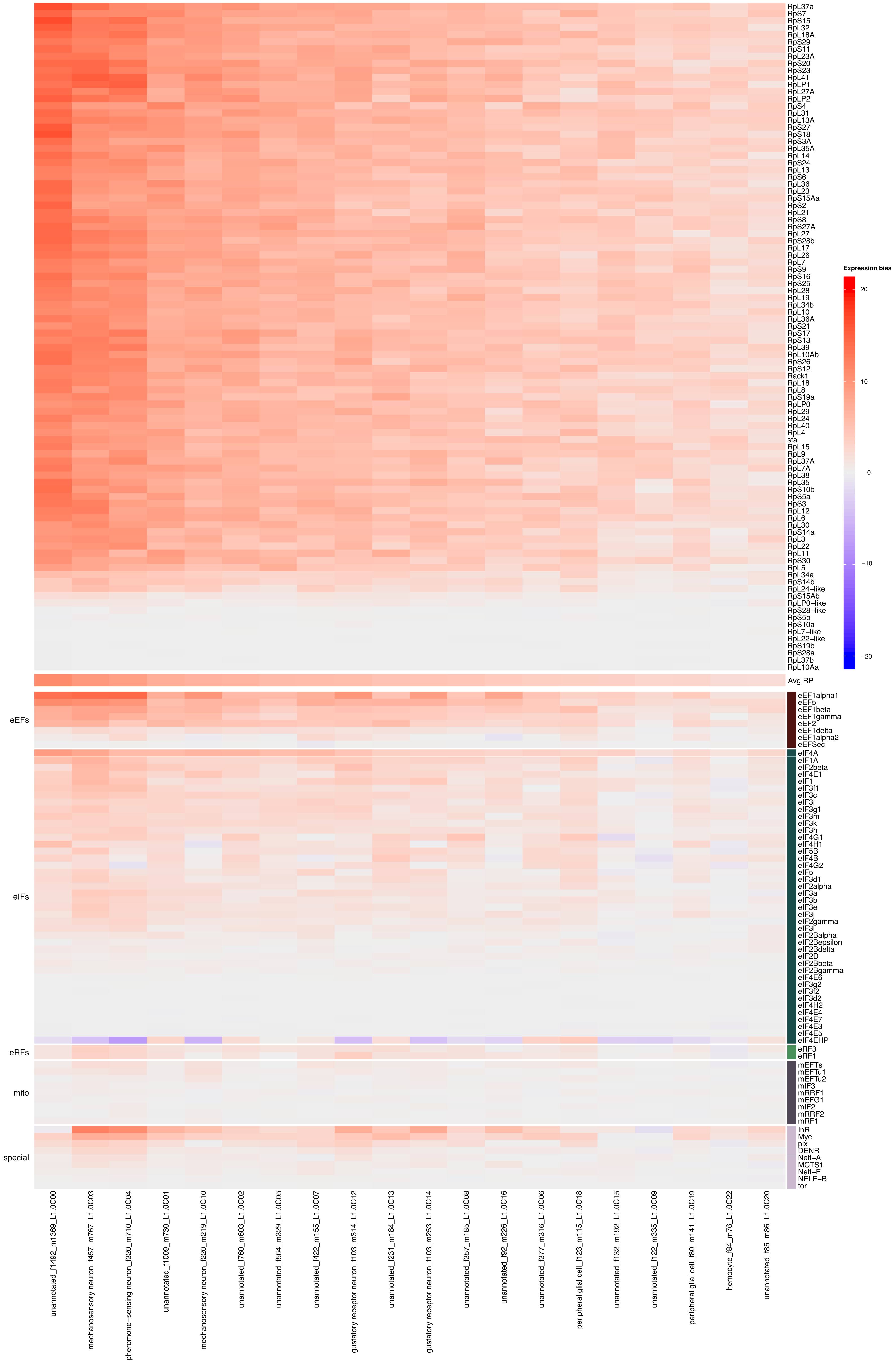

### Supp Fig S18

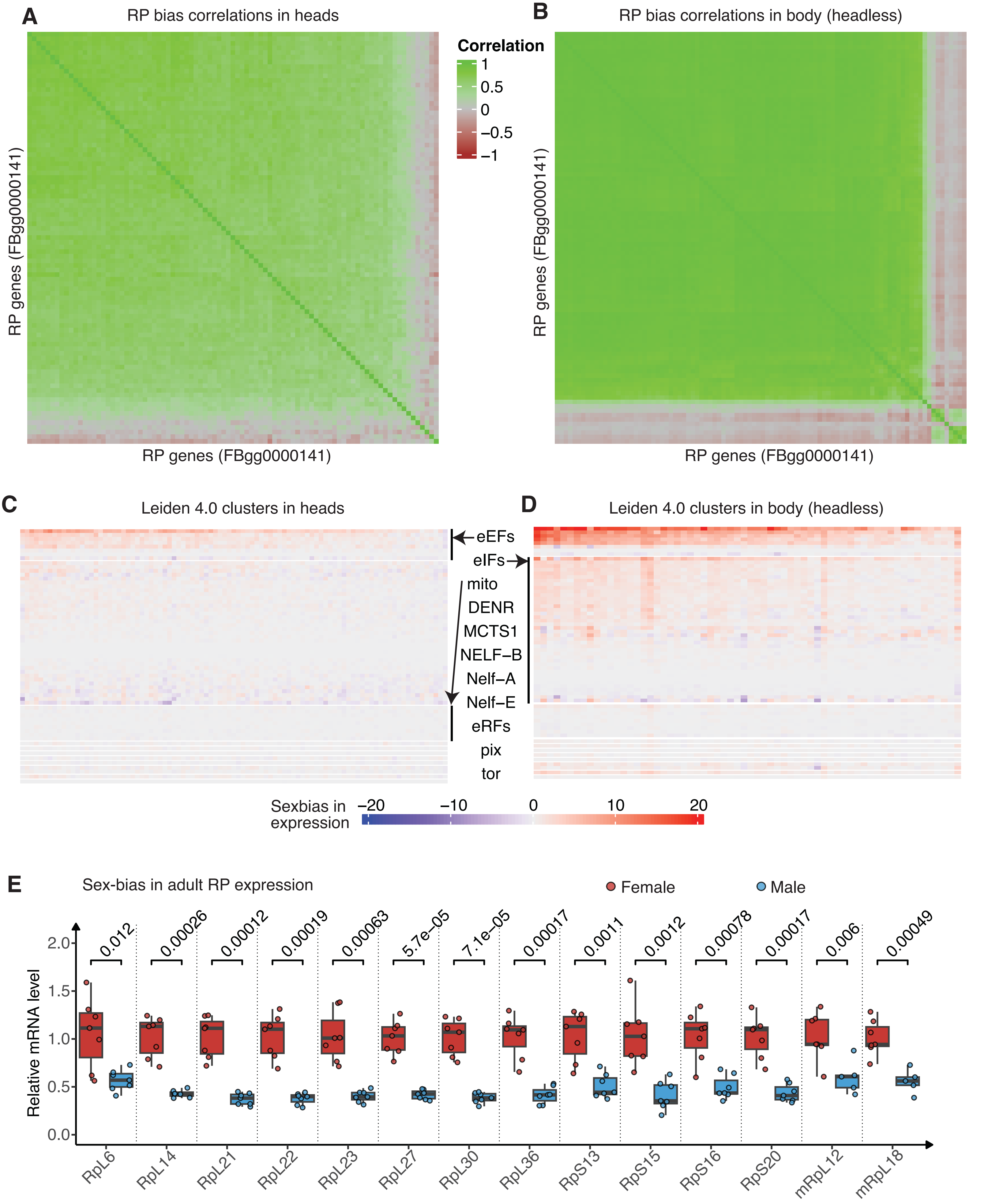

### Supp Fig S19

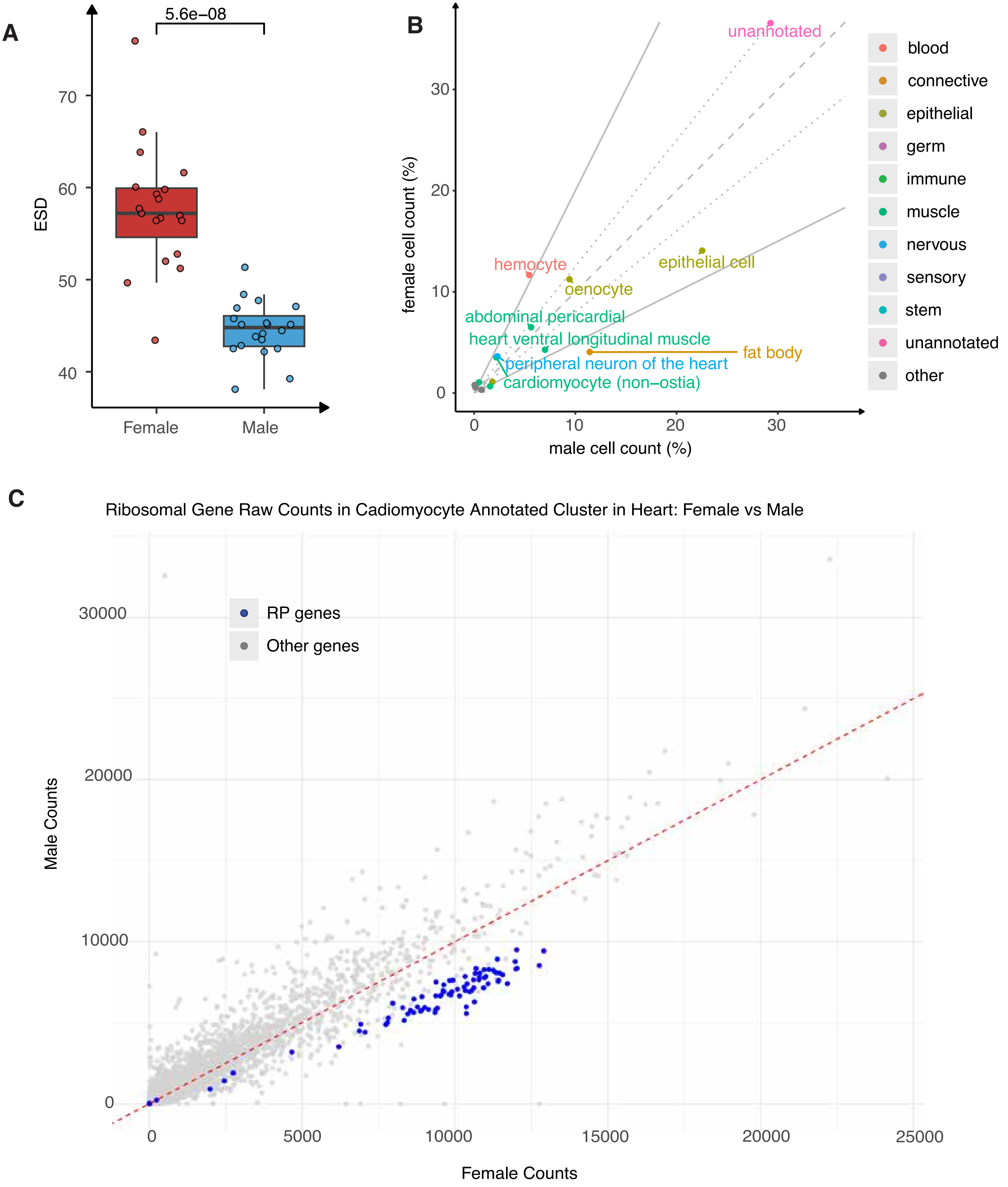

### Supp Fig S20

#### Figure S19

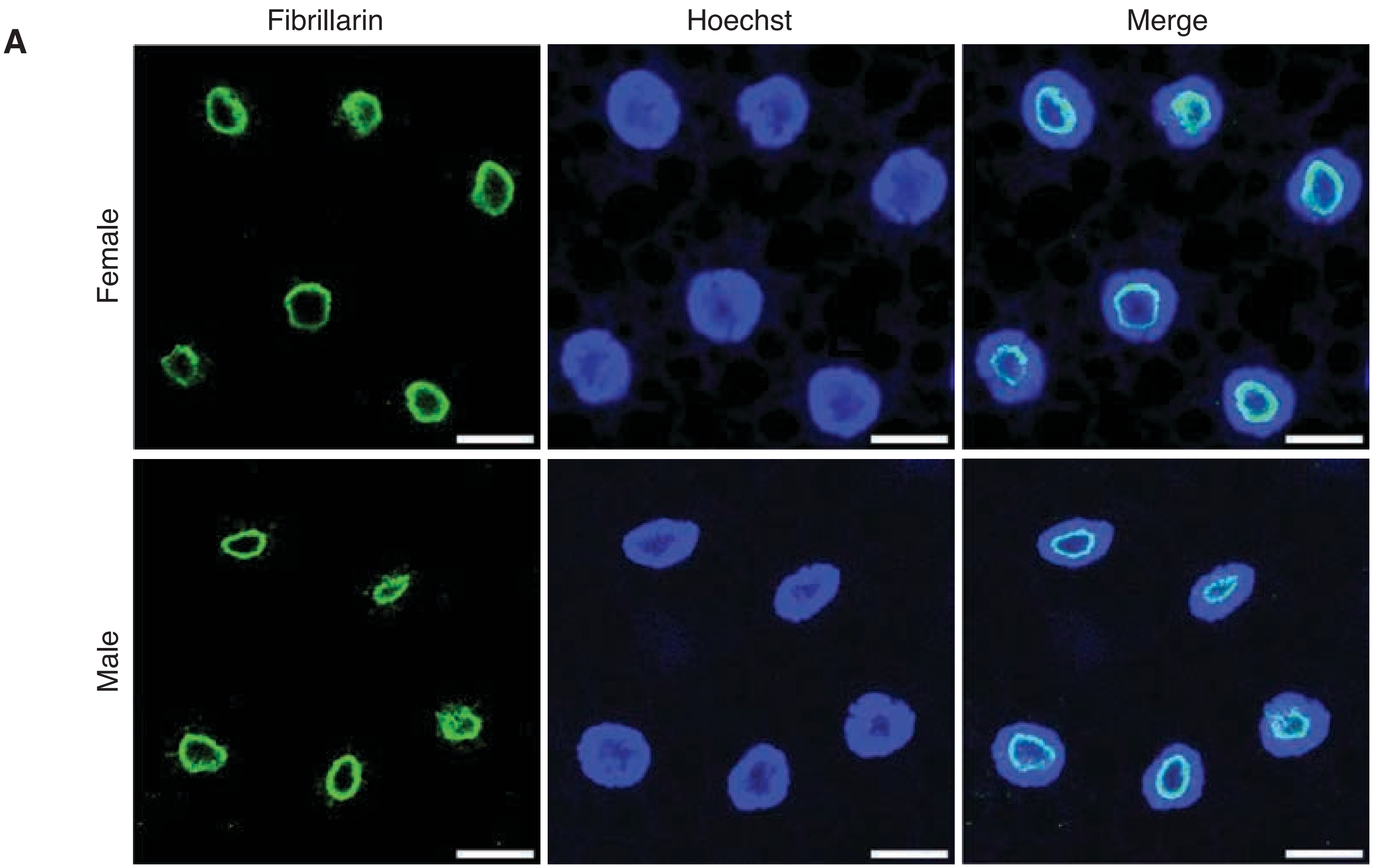
